# A super-pangenome of the North American wild grape species

**DOI:** 10.1101/2023.06.27.545624

**Authors:** Noé Cochetel, Andrea Minio, Andrea Guarracino, Jadran F. Garcia, Rosa Figueroa-Balderas, Mélanie Massonnet, Takao Kasuga, Jason Londo, Erik Garrison, Brandon Gaut, Dario Cantu

## Abstract

Capturing the genetic diversity of wild relatives is crucial for improving crops because wild species are valuable sources of agronomic traits that are essential to enhance the sustainability and adaptability of domesticated cultivars. Genetic diversity across a genus can be captured in super-pangenomes, which provide a framework for interpreting genomic variations. Here we report the sequencing, assembly, and annotation of nine wild North American grape genomes, which were phased and scaffolded at chromosome scale. We generate a reference-unbiased super-pangenome using pairwise whole-genome alignment methods, revealing the extent of the genomic diversity among wild grape species from sequence to gene level. The pangenome graph captures genomic variation between haplotypes within a species and across the different species, and it accurately assesses the similarity of hybrids to their parents. The species selected to build the pangenome are a great representation of the genus, as illustrated by capturing known allelic variants in the sex-determining region and for Pierce’s disease resistance loci. Using pangenome-wide association analysis, we demonstrate the utility of the super-pangenome by effectively mapping short-reads from genus-wide samples and identifying loci associated with salt tolerance in natural populations of grapes. This study highlights how a reference-unbiased super-pangenome can reveal the genetic basis of adaptive traits from wild relatives and accelerate crop breeding research.

## Introduction

The grapevine (*Vitis vinifera*) is an economically important crop of global significance, cultivated across the world to produce grape-based commodities such as wine, raisins, and grape juice. Historically, its productivity has been endangered by its high susceptibility to biotic and abiotic stresses, leading to the widespread practice of grafting *V. vinifera* cultivars on rootstocks derived from a handful of wild grape species [1,2]. The *Vitis* genus is composed of ∼70 interfertile species [3] with the North American wild species holding particular significance due to their extensive genetic diversity and historical contribution to the development of disease-resistant cultivars and rootstocks [2]. The geographical distribution of the grape species in North America contributes to their diverse array of stress tolerance traits (e.g., salt and drought tolerance, resistance to bacterial and fungal diseases, etc). Dioecious and interfertile, natural populations of wild grapes have evolved and adapted to the environmental conditions of their respective habitats and through interspecific introgression [4]. Their tolerance to biotic and abiotic stress enables them to thrive in a variety of climates and ecological niches, making them valuable genetic resources for breeding programs.

Although several grape genomes have been sequenced (e.g., [5–7]), working separately with distinct references impedes the comprehensive understanding of the species’ or genus’ genetic architecture. Numerous reports indicate that the use of a single reference genome directly impacts the effectiveness of identifying intraspecific genetic variations because reference mapping often fails to capture novel or highly divergent sequences within a species [8,9]. These limitations have motivated the development of pangenomes, which are transformative because they provide a comprehensive view of the genetic landscape within a species that may be otherwise overlooked [9]. The desire to capture an even broader spectrum of genetic diversity, encompassing wild relatives and diverse cultivars, further led to the generation of super-pangenomes, surpassing traditional pangenomes by incorporating genus-level diversity [8]. Super-pangenomes enable the identification of rare or unique genetic variations, population-specific alleles, and adaptive traits, providing a rich resource for crop improvement. Additionally, genus-level pangenomes can shed light on evolutionary history, domestication processes, and genetic relationships within a genus, contributing to a deeper comprehension of their genetic potential [8]. While pangenomes can have multiple definitions, graph-based methods are generally acknowledged to be the most comprehensive for eukaryotic genomes, because they include non-coding regions, regulatory elements, and intergenic regions, thereby enabling a more holistic understanding of genomic architecture and functional elements [10]. To date, however, super-pangenomes in plants have been limited by reducing diploid genomes to a haploid representation, by a constrained input order or phylogenetic dependencies, or by cataloging genetic variants against a single reference. None of these approaches overcome reference biases, and they often do not provide nucleotide-level resolution. Here we overcome these limitations by sequencing and assembling phased diploid genomes and building a super pangenome using all-vs-all whole-genome alignments. This approach is, by definition, reference-unbiased. It therefore overcomes a reliance on reference or tree-guided approaches, in an effort to better capture structural variations between distant species at the nucleotide-level resolution [11].

Here, we describe a reference-unbiased super-pangenome of North American wild grapes in the genus *Vitis* with as a main objective to identify and analyze the interspecific genetic variability. To build the super-pangenome, we first sequenced and assembled diploid, chromosome-scaled genomes from nine representative species: *V. acerifolia*, *V. aestivalis*, *V. arizonica*, *V. berlandieri*, *V. girdiana*, *V. monticola*, *V. mustangensis*, *V. riparia*, and *V. rupestris*. From these, we built a super-pangenome graph from all-vs-all chromosome alignments. From the generated graph, we inferred properties of the genic pangenome and extracted inter-species sequence variations. Our work has captured an unprecedented view of genetic diversity among North American wild grape species and effectively identified significant associations with salt tolerance using pangenome-wide association analysis (pan-GWAS).

## Results

### Assembly, annotation, and phasing of nine North American *Vitis* genomes

To construct a comprehensive representation of the North American *Vitis* genus, nine accessions were selected to be characteristic of their respective taxonomical group as defined previously based on their geographical distribution and phenological differences [3] and for their potential in breeding programs (Fig. 1a-j, Additional File 1: Table S1).

**Fig. 1:**
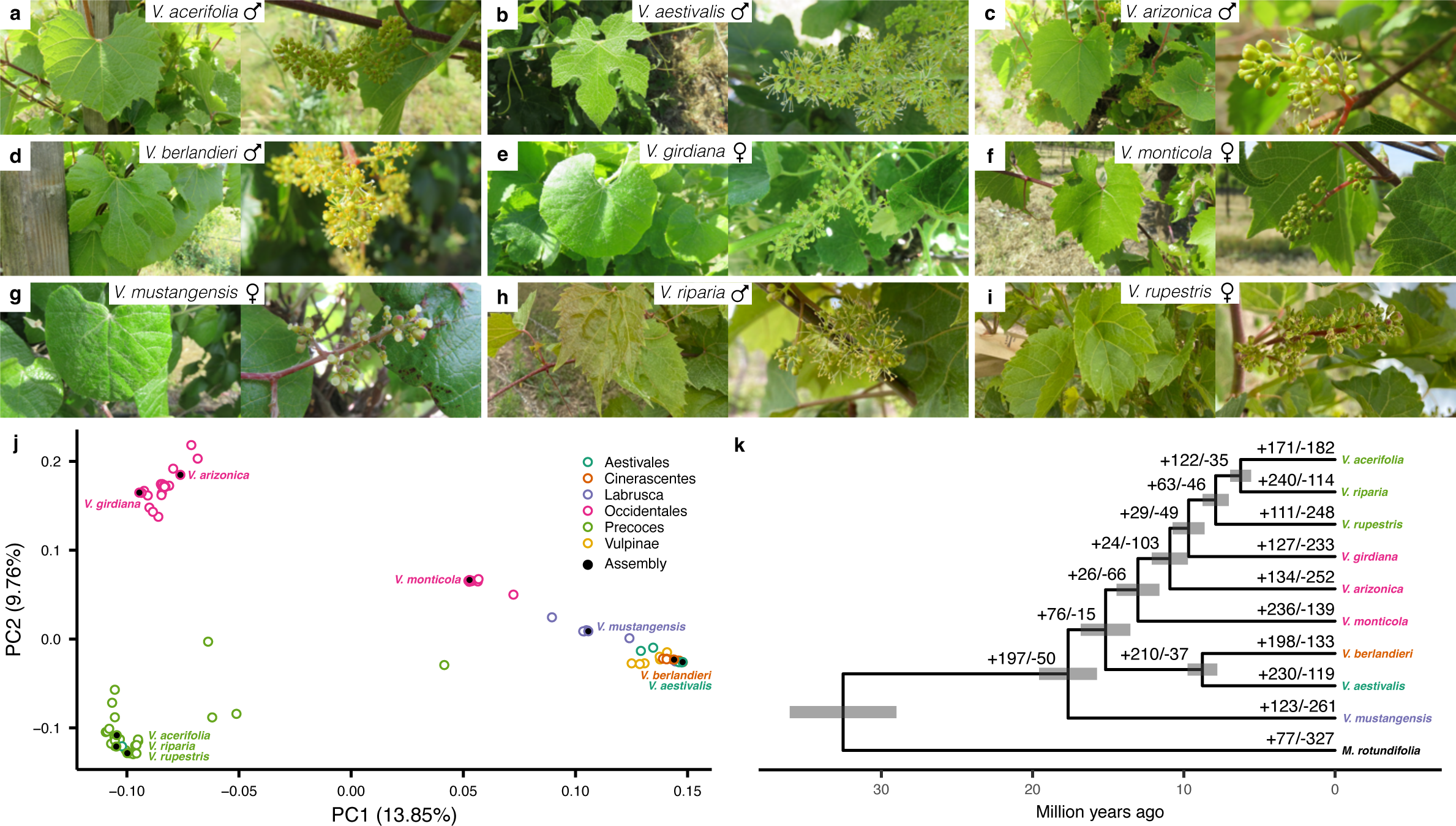
Selection of nine wild North American *Vitis* species. **a**, Pictures from a representative leaf (left) and flower (right) of *V. acerifolia* (**a)**, *V. aestivalis* (**b**), *V. arizonica* (**c)**, *V. berlandieri* (**d**), *V. girdiana* (**e**), *V. monticola* (**f)**, *V. mustangensis* (**g**), *V. riparia* (**h**), and *V. rupestris* (**i**), respectively. **j**, Principal component analysis results on SNP data derived from samples of natural populations (n = 90) aligned on *Vitis vinifera* Cabernet Sauvignon. Colors divide the *Vitis* genus into six groups. The accessions selected for assembly are represented by black-colored points. **k**, Phylogenetic relationships among the selected species. The phylogenetic tree was constructed based on 2,338 single-copy gene orthogroups and the branches represent divergence times in million years. *M. rotundifolia* was used as an outgroup to root the tree. Positive and negative numbers indicated on the branches represent expansions and contractions, respectively, among the 450 rapidly evolving orthogroups as determined by gene family evolution analysis.

Genomes were sequenced using single-molecule real-time sequencing (SMRT; Pacific Biosciences) with 114-255X depth (Additional File 1: Table S2). Optical mapping (Bionano) data were generated to produce consensus genome maps for seven of the genomes with 791-3,074 depth (Additional File 1: Table S3). The Falcon-Unzip contigs (N50 = 0.31-1.14 Mb; Table 1) were scaffolded using the genome maps to produce diploid hybrid assemblies (N50 = 1.13-9.22 Mb; Table 1). Protein-coding gene loci were annotated through an extensive *ab-initio* prediction pipeline relying on both Iso-Seq full-length transcripts and RNA sequencing (RNA-Seq) short reads (Additional File 1: Table S4). After the generation of consensus models, isoforms prediction, filtering, and functional annotation, 57,003-74,142 genes were annotated in the nine genomes (Table 1), covering on average 33% of the genome. Each species showed a BUSCO score exceeding 95% with over 50% duplication supporting the diploid representation of the assemblies.

**Table 1.** Statistics for genome assembly and annotation for the nine selected *Vitis* species.

| Species | Accession | Sequence length (Mb)<sup>1</sup> | Anchored sequence (%) | Contig N50 (Mb) | Scaffold N50 (Mb) | Gene count<sup>1</sup> | Anchored gene (%) | Repeats (%)<sup>1</sup> | BUSCO (%)<sup>1</sup> |
| --- | --- | --- | --- | --- | --- | --- | --- | --- | --- |
| *V. acerifolia* | 9018 | 486\|482\|55 | 94.6 | 0.31 | 1.74 | 31,605\|31,193\|2,751 | 95.8 | 44.7\|44.8\|56.7 | 90.8\|90.5\| 5.5 |
| *V. aestivalis* | T52 | 548\|545\|97 | 91.8 | 0.96 | 2.45 | 32,795\|33,222\|5,373 | 92.5 | 47.8\|47.9\|53.4 | 94.6\|93.8\|10.1 |
| *V. arizonica* | B40-14 | 466\|423\|66 | 93.1 | 0.68 | 9.21 | 28,043\|25,925\|3,035 | 94.7 | 47.3\|47.0\|59.8 | 89.9\|83.9\| 6.1 |
| *V. berlandieri* | 9031 | 554\|549\|114 | 90.6 | 1.06 | 2.68 | 33,866\|33,472\|6,804 | 90.8 | 47.7\|47.5\|52.1 | 95.3\|95.0\|12.6 |
| *V. girdiana* | SC2 | 464\|476\|65 | 93.5 | 1.14 | 4.08 | 28,765\|29,882\|2,884 | 95.3 | 47.2\|46.4\|58.9 | 92.9\|81.7\| 5.6 |
| *V. monticola* | TX6703 | 521\|505\|99 | 91.2 | 0.57 | - | 33,281\|32,299\|5,941 | 91.7 | 45.9\|45.3\|51.3 | 93.7\|88.2\|16.9 |
| *V. mustangensis* | C114-96 | 454\|432\|71 | 92.6 | 0.49 | 1.13 | 27,803\|26,177\|3,510 | 93.9 | 44.5\|44.3\|56.3 | 92.8\|85.3\|11.0 |
| *V. riparia* | HP-1 | 520\|506\|113 | 90.1 | 1.09 | - | 32,717\|31,793\|6,778 | 90.5 | 46.9\|46.8\|52.6 | 95.9\|91.4\|15.5 |
| *V. rupestris* | B38 | 449\|448\|41 | 95.6 | 0.80 | 9.22 | 29,038\|28,798\|1,621 | 97.3 | 45.4\|45.3\|58.8 | 93.2\|92.6\| 4.6 |

<sup>1</sup> The values represent haplotype 1, haplotype 2, and unplaced sequences.

The tool HaploSync [12] coupled with a high-density *Vitis* genetic map [13] was used to construct nine diploid sets of nineteen chromosomes with a target of at least 90% of anchored sequences and protein-coding gene loci (Table 1). For each genome, the markers and the relation primary contigs/haplotigs were used as constraints to simultaneously phase the sequences into two haplotypes and attribute them to a chromosome. The phasing correctness was evaluated at every step of the scaffolding using both the allelic gene content and DNA sequencing (DNA-Seq) short-read coverage (Additional File 1: Table S5) to ensure the accuracy of the diploid phasing and prevent scaffolding issues such as haplotype switches. Gap filling was performed with the remaining unplaced sequences based on the pairwise relationship between the haplotypes. The best assembly performance reached 95.6% of sequence anchorage to the chromosomes for *V. rupestris*, representing 97.3% of its gene content. For all the genomes, the unplaced sequences were composed of short sequences with an N50 = 40,653-63,859 bp (Additional File 1: Table S6), mostly enriched in repeats (Table 1). To estimate the levels of heterozygosity in the nine *Vitis* genomes, we compared the assemblies of their phased haplotypes and considered every type of variants and any size. As expected, each genome presented high levels of variation between haplotypes, ranging from 16.79 to 23.77% of the genome sequence impacted by variants when the two corresponding haplotypes were compared (Additional File 1: Table S7) falling in the range of previous conservative estimates such as in Chardonnay (15.3%) [14] or Pinot Noir (11.2%) [15]. SNPs and INDELs represented almost the entirety of the variants detected (99% on average), while translocations or inversions were very few. SNP was the most abundant type of variant with 1 SNP every 238-1159 bp (Additional File 2: Figure S1) as reported previously for wild grapes species [16,17]. INDELs were more spread with an average distribution six times wider than for the SNPs (1 INDEL every 1652-5030 bp) (Additional File 2: Figure S1).

Despite such variations at the sequence level, gene content was well conserved between haplotypes, with genic-hemizygosity rates ranging from 2.09 to 4.95%, except for *V. arizonica* (9.35%) (Additional File 1: Table S8). The phasing integrity was further supported by the BUSCO scores at 93.2 ± 2.0% for haplotype 1 and 89.2 ± 4.7% for haplotype 2 (Table 1). The complete diploid representation of the genome assemblies was further supported by their length; representing approximately twice the haploid sizes estimated by flow cytometry, but also by the presence of 1.90-2.27X the haploid gene content of PN40024 [5] (Additional File 1: Table S9). Altogether, the assembly of these nine high-quality genomes provided the foundation to build a comprehensive super-pangenome using comparable assembly/scaffolding qualities. Orthogroups were identified using the haplotype 1 as representative as it presented systematically the highest BUSCO score of the two haplotypes and was generally the longest haplotype with the most genes. Given the nine reference genomes, we aligned in a pairwise configuration the nine haplotype 1 proteomes, resulting in the identification of 2,338 single-copy orthogroups (SCO). We gathered the SCO alignments into a supermatrix and used it to produce a species-wide phylogeny that matched previous inferences [18,19] and separated species by known *Vitis* groups [3] (Fig. 1k). We also estimated divergence times at each node using molecular clock method [20] (Fig. 1k). Finally, after the selection of one representative isoform per gene, we used the same proteomes to evaluate gene families across the North American *Vitis* phylogeny, identifying 450 rapidly evolving orthogroups (Fig. 1k). Interestingly, the gene families involved in the contraction/expansion events were significantly enriched for gene ontology (GO) terms such as response to stress, signaling, response to stimulus, and cell communication, suggesting that they might be involved in the adaptation of each species (Additional File 1: Table S10).

### A nucleotide-level super-pangenome graph for the North American *Vitis* genus

The 18 haplotype genomes (9 genomes x 2 haplotypes) were considered as distinct inputs for the super-pangenome construction. After pairwise comparisons of assemblies between species, we inferred that 27.10 to 34.20% of the genome from each species was affected by genetic variants (Additional File 1: Table S11), values that greatly exceeded heterozygosity (16.79-23.77%). We used nucleotide variants to estimate an average rate of 2.4×10^-8^ ± 6×10^-9^ variants per nucleotide per year between *Vitis* genomes using the divergence times inferred in the clock-calibrated phylogeny (Fig. 1k). To avoid any bias towards a specific reference, the pipeline to construct the pangenome relied on all pairwise sequence alignments of the 18 genomes [21]. The workflow followed the PanGenome Graph Builder (PGGB) pipeline [11] which involves three main steps; wfmash to generate all the pairwise alignments, ii) seqwish to induce the pangenome graph from the alignments, and iii) smoothxg to polish the pangenome graph. In the resulting graph topology, genomic sequences are represented as “nodes” that can be as small as one base pair. The chromosomes are stored as “paths” which are a linear representation of their DNA as traversals through the nodes.

The generated pangenome graph had a compression ratio per chromosome of 81% on average (Fig. 2a) and contained 342 paths, each representing a chromosome of the input haplotype genomes. The graph structure was composed of ∼ 200 million nodes (Additional File 2: Figure S2) for a total length of ∼ 1.7 Gbp (Fig. 2b) and representing ∼1.5X the size of the diploid genome for a single *Vitis* species. Nodes were categorized into three different classes according to the number of species containing them: i) core genome; nodes present in the nine species, ii) dispensable genome; nodes present in more than one species but not all, and iii) private genome; genome-specific nodes. The nodes in the private genome covered the largest portion of the pangenome (1.1 Gb). However, most of the nodes (> 100 million) were in the dispensable genome while both the core and the private genomes were composed of about 50 million nodes each (Additional File 2: Figure S2). At the haplotype level, the core genome represented about half the genome size (48% on average), while the dispensable and the private genomes contained on average 36% and 16% of the genome, respectively (Fig. 2c).

**Fig. 2:**
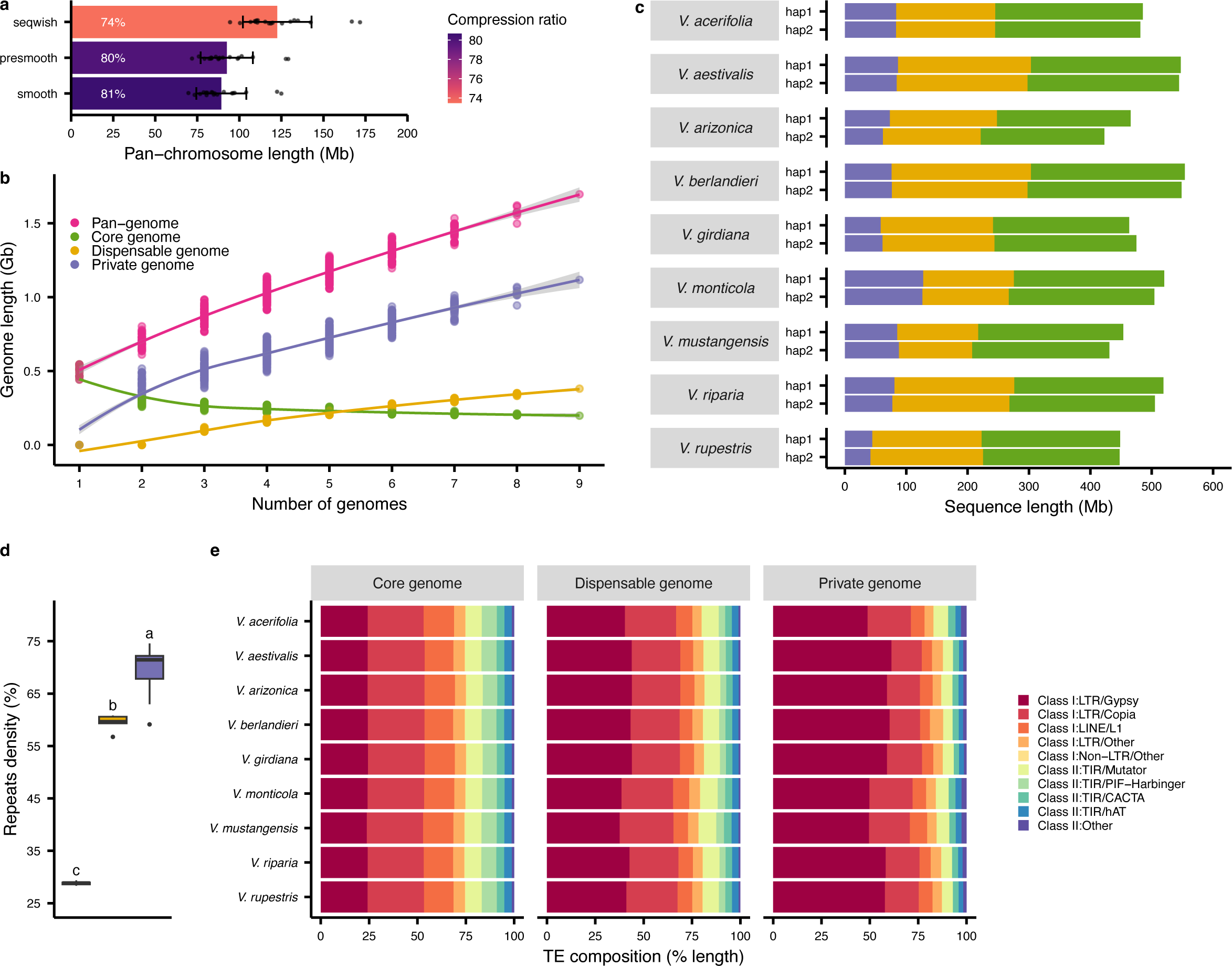
Properties of the nucleotide-level super-pangenome graph. **a**, Compression ratio of the pangenome. For each step of the pangenome construction, raw (seqwish) and post-polishing (smoothxg), the mean ± sd (n = 19 chromosomes) of the total length of the unique nodes representing each chromosome is represented. The compression is expressed as a ratio of the total size of a chromosome in the graph compared with the sum of the lengths of the same chromosome in the 18 haplotypes used as input. **b**, Graph-based pangenome modeling. For every combination of 1-9 genomes, the total length of the unique nodes is represented per class. The line represents smoothed conditional means with a 0.95 confidence interval. **c**, Pangenome class composition per haplotype for each genome. **d**, Repeat density observed in the different node classes of the pangenome. Significant groups were determined using a multiple comparison test after Kruskal-Wallis with a *P* value < 0.05. The middle bars represent the median while the bottom and top of each box represent the 25^th^ and 75^th^ percentiles, respectively. The whiskers extend to 1.5 times the interquartile range and data beyond the end of the whiskers are plotted individually as outlying points. Panels **b**, **c**, and **d** share the same color legend. **e**, Transposable elements composition in the different pangenome classes. Categories with unspecified repeats or representing < 2% of the repeat content were classified as “Other”.

One interesting feature of these analyses was that the private genome was composed of a significantly higher proportion of repeats compared to the other classes (Kruskal-Wallis, *P* value < 0.05). The private genome was ∼ 40% more repetitive than the core genome (Fig. 2d). Class I transposable elements (TEs) represented the majority (80.3%) of the TEs in the pangenome (Additional File 2: Figure S3). Remarkably, 56% of the private TEs were long terminal repeats (LTR) *Gypsy* retrotransposons, which represented only 24% of the core TEs (Fig. 2e). The over-representation of Gypsy elements in the private genome suggests they are strongly associated with genome divergence between species and may play a pivotal role in grape genome evolution.

### Intra and inter-specific relatedness is captured in the pangenome

To evaluate the sensitivity of our pangenome, we added the previously published diploid genomes [7] of three hybrids to the pangenome, all derived from the cross of North American species included in our pangenome: 101-14 Millardet et de Grasset (101-14 Mgt: *V. riparia* x *V. rupestris*), Richter 110 (110R: *V. berlandieri* x *V. rupestris*), and Kober 5BB (*V. berlandieri* x *V. riparia*). As expected, the addition of the three hybrids with the nine previous genomes led to a decrease in the relative private and core genome sizes in favor of a larger dispensable genome, both in terms of length and node number (Additional File 2: Figure S4). We incorporated these hybrid genomes to evaluate whether we could accurately assess their origins and to characterize their structural effects on the pangenome graph topology. We established different types of comparisons to assess whether the pangenome graph was able to reflect correctly different degrees of similarity between close or distant species.

To evaluate genome relatedness, we focused on the three hybrids and the three parental species from which the hybrids were derived. We quantified the degree of relationship between the different genomes using the total length of the nodes they shared as a metric (Fig. 3). Three levels of comparisons were investigated: intra-genome by the comparison of the two haplotypes within each genome (Fig. 3a-c), inter-haplotype by the comparison of the hybrids’ haplotypes to their parents (Fig. 3d-f), and inter-genome comparing haplotypes of different species (Fig. 3f-i). First, intra-genomic comparisons were performed between the two haplotypes of each genome; on average, 60.2% was shared between the two haplotype sequences of each species (Fig. 3b-c). Notably, the parents presented a higher percentage of shared node space between their haplotypes (65.9% on average) compared with the hybrids (54.4% on average). The hybrids having *V. berlandieri* as a parental species showed the lowest percentage (Richter 110R: 51.7%; Kobber 5BB: 50.9%), probably because this species is genetically distant from the two other parents *V. riparia* and *V. rupestris* (Fig. 1k). Secondly, we evaluated if the pangenome could detect the relatedness of the hybrids with their parental species. Each hybrid haplotype was derived from a different species, as determined during their assembly [7]. When we compared each hybrid haplotype (e.g. 110R haplotype berlandieri annotated 110R be on Fig. 3d) with the haplotypes of its parental genome (i.e. *V. berlandieri* haplotype 1 or 2 annotated be hap1 or hap2 on Fig. 3d), we found on average 58.2% of their nodes being shared. *V. rupestris* showed the most conserved node space with its corresponding hybrids 110R and 101-14 Mgt (Fig. 3e-f). Lastly, the smallest percentages of shared node space were detected when the genomes were compared in an inter-specific configuration (Fig. 3g-i). As expected, *V. riparia* and *V. rupestris* haplotypes showed the highest shared node space since there are closer to each other than *V. berlandieri*. Despite those similarities, the overall shared node content between different species remained lower than the percentages observed when hybrid haplotypes were compared with their parental species. Altogether, these results illustrate the importance of haplotype-resolved genomes, demonstrate that the super-pangenome graph captures both intra and inter-specific variants, and accurately represents the relationship between parental species and hybrids, thus reinforcing the high sensitivity of the super-pangenome.

**Fig. 3:**
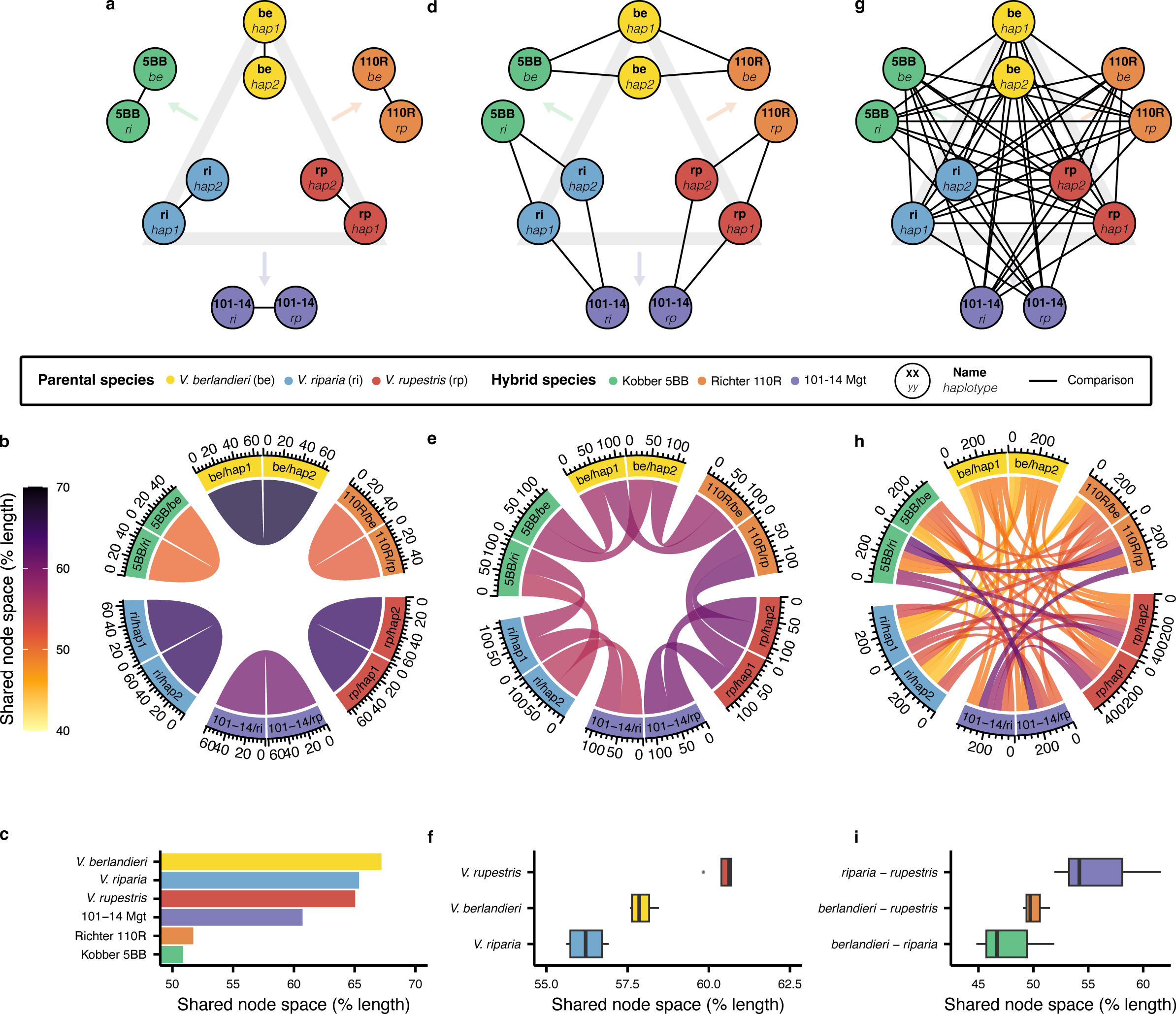
Graph sensitivity assessment using phased hybrids and their parental species. On the top row, the base diagram represents the relation between the parents and their progeny. The grey triangle contains the three parental species used to give the three hybrids indicated after each arrow. The black lines connect the haplotypes that were compared. Three different types of comparison are represented: on the left (**a**-**c**), haplotypes are compared within species; in the middle (**d**-**f**), haplotypes are compared between the hybrids and their parents; on the right (**g**-**i**), the haplotypes are compared in an interspecific manner. The middle row represents the individual share node space for each comparison, the percentage values being indicated as a color gradient. The bottom row contains a summarized version of the chord diagrams represented either as a barplot for single values (**c**; n = 1) and boxplot when multiple values are involved (**f**; n = 4 and **i**; n = 15). The middle bars represent the median while the bottom and top of each box represent the 25^th^ and 75^th^ percentiles, respectively. The whiskers extend to 1.5 times the interquartile range and data beyond the end of the whiskers are plotted individually as outlying points.

### Gene-based pangenome inferred from the pangenome graph

We used the super-pangenome to study the distribution and characteristics of genes among the nine *Vitis* species. Genes were categorized into three different genome classes according to their node composition: i) core genes, when the gene sequence was composed of >80% of core nodes, dispensable genes, when the sequence was composed of >80% dispensable nodes, and iii) private genes when the sequence was composed of >80% private nodes. The genes not falling in any of the above categories were classified as ambiguous; these represented on average only 2.6% of the gene content and were excluded from subsequent analyses (Additional File 1: Table S12). We examined the core, dispensable, and private categories as a function of species sampling: as genomes were added to the pangenome, the number of private genes continuously increased, inflating the overall gene pangenome size (Fig. 4a). In contrast, the number of core genes tended to slowly stabilize around 40,000 genes (Fig. 4a, Additional File 1: Table S13), representing more than 67% of gene content per species on average (Fig. 4b, Additional File 1: Table S13). Although the number of private genes continually increased with the addition of species, private genes represented were less numerous than core genes ranging from 2,580 to 9,145 genes per species.

**Fig. 4:**
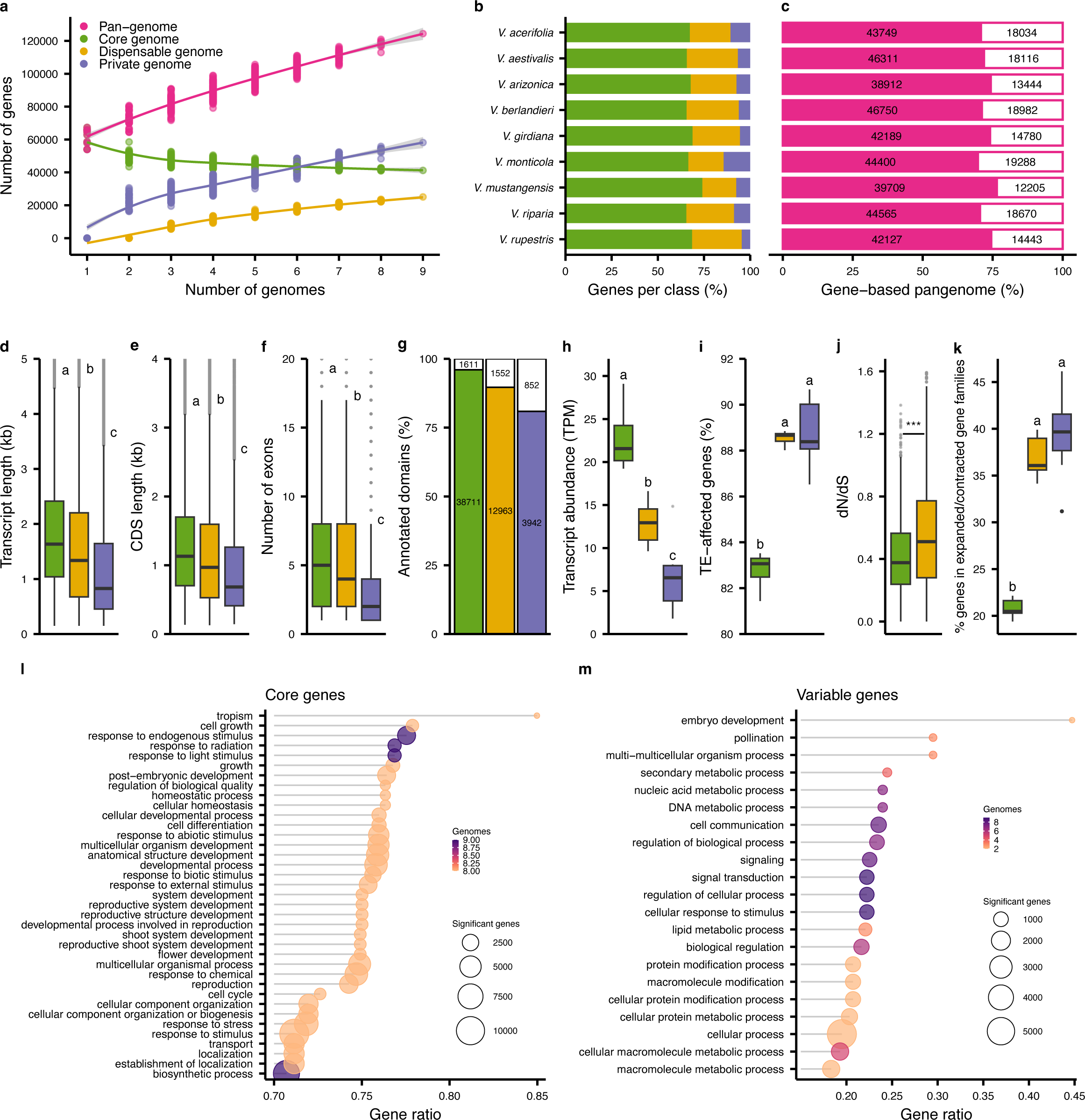
Characterization of genes in the super-pangenome. **a**, Gene-based pangenome modeling. For each combination of 1-9 genomes, the number of genes is represented per class. The lines represent smoothed conditional means with a 0.95 confidence interval. **b**, Gene class composition for each genome. **c**, Percentage of genes attributed to the same class in both the graph-derived and orthology-based pangenomes. The number of consistent genes between both approaches is represented on the left of the bar while the different are on the right. **d**, Transcript length (kb); **e**, CDS length (kb); **f**, Number of exons; **g**, Percentage of annotated domains; **h**, Transcript abundance (TPM); **i,** TE-affected genes; **j,** dN/dS ratios, and **k**, Expanded/Contracted gene families per class of genomes. *P* values were determined using ANOVA and significant groups were assigned following Tukey’s “Honest Significant Difference” method. The middle bars represent the median while the bottom and top of each box represent the 25^th^ and 75^th^ percentiles, respectively. The whiskers extend to 1.5 times the interquartile range and data beyond the end of the whiskers are plotted individually as outlying points. For the dN/dS ratios (**j**), *P* values were determined using a two-tailed Student’s t-test. To prevent the compaction of the y-axis by extreme outlying points, the upper limit was capped. Panels **a**-**k** share the same color legend. Enriched gene ontology terms for the core genes (**l**) and the variable (dispensable + private) genes (**m**) are represented as circles with their size depending on the number of genes involved and their color based on the number of genomes having the term detected as significant. For each GO term, the number of significant genes relative to the total number of annotated genes for the term is represented as a gene ratio on the x-axis.

These results were supported by a classic orthology analysis based on colinearity. On average, 72.6 ± 2.3% of the genes were consistently categorized as core, dispensable, or private genes between the graph-inferred and the orthology-based gene pangenomes (Fig. 4c, Additional File 1: Table S14). The core genome was the most conserved class, with 85.8 ± 1.76% consistency. The three classes of genes also exhibited remarkable structural differences. Dispensable and private genes were significantly shorter than the core genes, with correspondingly smaller transcript and CDS lengths and fewer exons (Fig. 4d-f). The longer core genes had more annotated domains (Fig. 4g). A striking association was also observed between gene expression and gene conservation within the genus. Transcript abundance was significantly higher for core genes, suggesting functional conservation relative to dispensable and private gene sets (Fig. 4h). The variable (i.e., non-core) genes had higher TE density around them (Fig. 4i), which may suggest that TEs contribute to their lower expression.

We also investigated the evolutionary properties of genes. For example, we calculated non-synonymous/synonymous substitution (dN/dS) ratios (Fig. 4j) for each gene class. On average, dispensable genes had higher average dN/dS values, reflecting higher sequence conservation among core genes. Among the genes belonging to the rapidly evolving families identified during the phylogeny analysis, the proportion classified as variable genes (dispensable/private) was twice as high (Fig. 4k) as the core genes (Kruskal-Wallis, *P* value < 0.05). This high proportion further suggests the possibility that dispensable genes contribute to the local adaptation of each species (Additional File 1: Table S15). Gene ontology (GO) enrichments supported this conjecture. For example, core genes were mainly associated with essential developmental functions such as the shoot system, flower development, and responses to stimulus. In contrast, variable genes were enriched in metabolic processes, cellular processes, and signaling (Fig. 4l,m). The enrichment of basic functions for the variable genes such as embryo development does not rule out the evolution of important functions for specific species. We focused particularly on nucleotide-binding site leucine-rich repeat (NLR) genes because they have been heavily studied in grapes for their association with disease resistance. Interestingly, NLRs possessing a toll/interleukin-1 receptor-like (TIR) domain, the TIR subclasses such as TIR-NBS-LRR genes, tended to be more abundant in the dispensable genome (Additional File 2: Figure S5). Our results consistently supported the strong, conservative selection of core genes, presumably due to their essential functions. The pool of private and dispensable genes does not seem to be conserved, at least under certain environmental conditions, suggesting that they might provide benefits for local adaptation to specific abiotic and/or biotic stresses and could be responsible for significant phenotypic variation among species.

### Structural variants derived from the pangenome graph topology

By definition, the pangenome graph embeds polymorphisms and structural variants for all the pairs of genomes used as input. To extract this level of information from the graph, we employed the module “deconstruct” from the tool vg v1.48.0 [22], selecting iteratively each genome in the graph as a reference. A variant analysis performed independently between genomes using NUCmer [23] provided similar results; the pangenome and NUCmer-based inferences were consistent presenting 90% and 76% of SNPs and INDELs, respectively (Additional File 1: Table S16). SNPs were the most abundant variant type within individual genomes (Fig. 5a). By classifying the variants in different groups of sizes, we observed that more than 96% of the total were short-length variants with 1-bp variants (most SNPs) as the most abundant size and the remaining set being shorter than 10 bp (Fig. 5b). For each species, 1 heterozygous SNP was detected every 239-888 bp between haplotypes representing 0.11-0.42% of their assembly (Additional File 1: Table S17) which is consistent with our independent pairwise comparison using NUCmer and reports from literature [16,17].

**Fig. 5:**
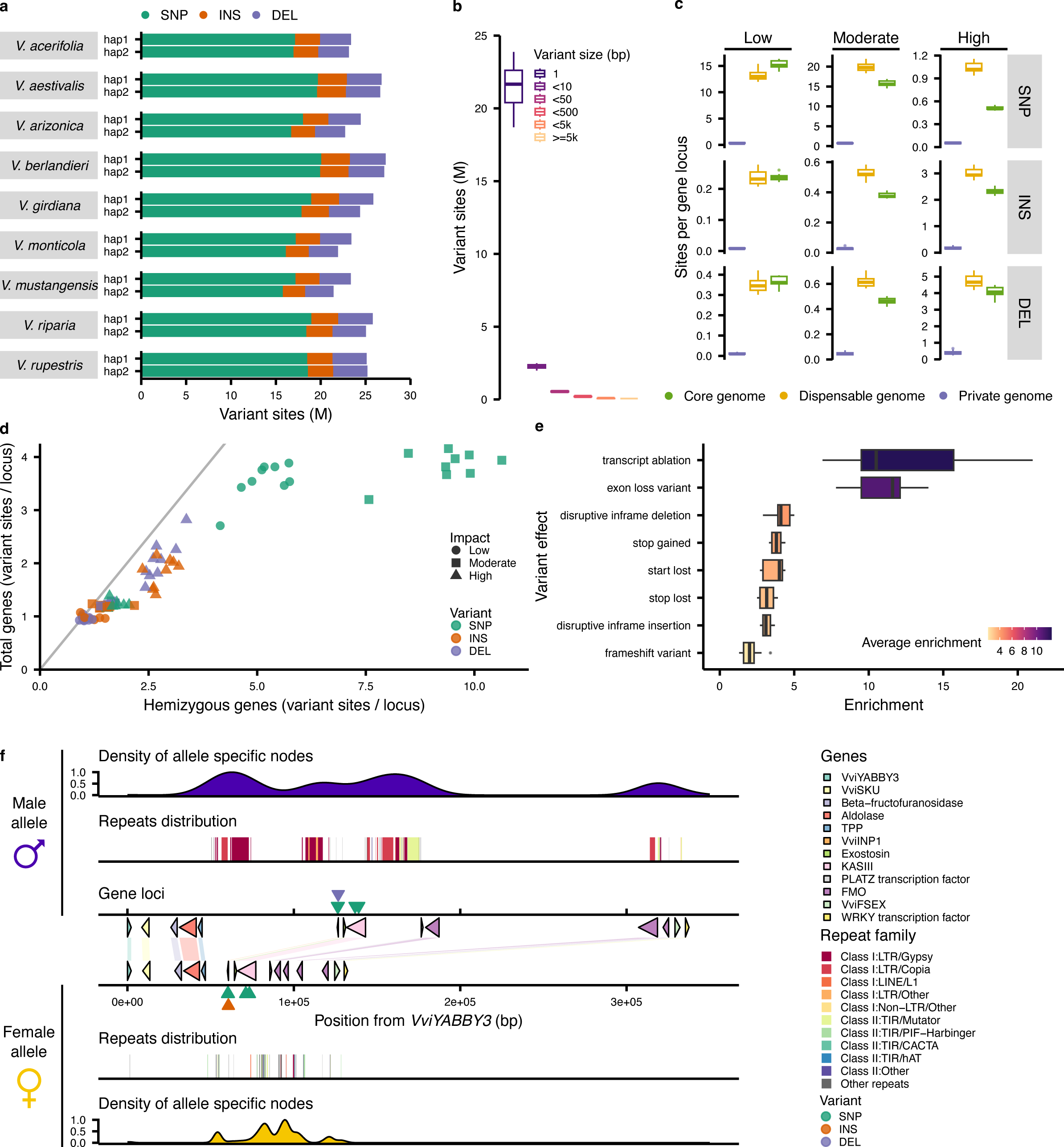
Genetic variants from the graph topology. **a**, Number of variant sites in each haplotype per genome. SNP = Single Nucleotide Polymorphisms, INS = Insertions, DEL = Deletions, M = Millions. **b**, Distribution of variants by size. **c**, Variant sites distribution per variant types and impacts for each pangenome class (n = 18 haplotypes). **d**, Number of variant sites per gene locus in hemizygous genes (x-axis) and all the genes impacted by variants (y-axis) (n = 9 genomes). **e**, Enrichment analysis results for the per variant effect identified in the hemizygous genes when compared with all the genes impacted by variants. Boxes are colored with the average enrichment values (n = 9 genomes). For the boxplot representations in **c** and **e**, the middle bars represent the median while the bottom and top of each box represent the 25^th^ and 75^th^ percentiles, respectively. The whiskers extend to 1.5 times the interquartile range and data beyond the end of the whiskers are plotted individually as outlying points. **f**, Pangenome features at the sex-determining region. *V. arizonica* B40-14 was used as representative for the region. For each allele (female and male), the density of allele-specific nodes, the repeats distribution, and the gene loci with the conserved variants are represented. The region is a sub-selection of the SDR spanning genes encoding VviYABBY3 to the WRKY transcription factor.

We annotated the variants using snpEff v5.1 [24] to estimate their impact on the protein sequence. They were classified in four categories of impact: low, mostly harmless to the protein; moderate, can affect protein effectiveness; high, disruptive impact in the protein; and modifier, usually non-coding variants. Generally, low-impact variants tended to be similarly distributed between the core and the dispensable genomes (Fig. 5c). When the impact on the protein sequence was moderate or high, most of the variants were found in the dispensable genome rather than in the core genome (Fig. 5c). The most striking evidence was illustrated by the SNPs. While they were similarly abundant in both the core and the dispensable genomes for at low impact, variants with moderate impact were more abundant in the dispensable genome. The difference was even larger for the number of high impact variant with ∼2x more variants in the dispensable genome (Fig. 5c). The private genome was particularly less impacted by variants when compared with the core or dispensable genomes confirming that the sequence of a private gene is mostly genome-specific and do not share many nodes (considered here as potential variant sites), if any, with the other genomes. Regarding the modifier sites (usually variants in non-coding regions), they were the most abundant type of impact and tend to display a similar distribution than the moderate variants (Additional File 2: Figure S6). Within the genes impacted by the variants, we decided to focus on the hemizygous gene sets we previously identified. We observed an overall higher abundance of variants in these gene loci (Fig. 5d) with more than 2x more sites on average for SNPs with moderate impact. Among the effects with high impact, transcript ablation and exon loss variant were the most enriched in the hemizygous gene loci compared with the total content of genes impacted by variants (Fig. 5e).

To validate the variants derived from the pangenome graph topology, we investigated its structure at the grape sex-determining region (SDR) (Fig. 5f). For both alleles (female and male), we observed a striking allele-specific accumulation between the *TREHALOSE-6-PHOSPHATE-PHOSPHATASE* (*TPP*) and *INAPERTURE POLLEN 1* (*VviINP1*) genes which correspond to the peak of linkage disequilibrium previously observed in this region [6]. In the females, allele-specific nodes were also accumulated around the Flavin-containing monooxygenase (FMO) loci which might be involved in the conservation of their function as they are not all protein-coding in the males [6]. The density of allele-specific nodes was also noticeably overlapping the repeats distribution suggesting an important role of repetitive elements in the larger size of the male allele. Few SNPs and INDELs were found to be conserved in an allele-specific manner. We focused on the variants impacting *VviINP1*, known to be involved in pollen aperture formation [6]. In the female allele, the sequence of this gene contains an 8-bp deletion which has been suggested to be responsible to the non-functionality of the corresponding protein [6]. Remarkably, the super-pangenome graph captured the known variants distinguishing the *INP1* male and the female alleles (Fig. 5f), including an allele-specific SNP at 60 bp from the *VviINP1* transcriptional start site (TSS) and the 8-bp INDEL starting at 74 bp from the TSS (Additional File 2: Figure S7). Moreover, the flower sexes of all the accessions used to build the pangenome were predicted correctly by their alleles in the pangenome and confirmed with phenotyping data from the field (Fig. 1a-i).

To further explore the variants embedded in the pangenome graph and gather information over multiple chromosomes, we also investigated polymorphisms associated with Pierce’s disease resistance. Eight peaks of association were previously detected in *V. arizonica* [25] (one of the input genomes integrated into the pangenome) using genome-wide association studies (GWAS). As the other genomes included in the pangenome were not selected for Pierce’s disease resistance, we expect to detect the polymorphisms. The nodes of the pangenome graph corresponding to those regions were extracted (Additional File 2: Figure S8). For 7 of the 8 association peaks, SNPs were detected in nodes that were either considered as variants from the graph topology or in the variable genome (private or dispensable). In other words, almost the entirety of the regions previously detected *de novo* by variant calling was represented in the pangenome graph structure. As a result, mapping the reads from the previous GWAS on the pangenome should in theory only require genotyping the embedded variants to detect the associated polymorphisms, thereby bypassing any variant calling.

### Pan-GWAS revealed loci associated with salt tolerance

In a previous study, 325 accessions from natural populations of 14 *Vitis* species were collected and evaluated for chloride exclusion, a process that confers salt tolerance in grapes [26]. Among the accessions used to build the pangenome, five were included in this earlier work and presented pronounced differences regarding salt accumulation in the roots and the leaves. In particular, *V. girdiana* accession SC2 was a high salt excluder. We sequenced the DNA of 153 samples from 12 species of the collection panel to perform a pangenome-wide association analysis (pan-GWAS). The variants extracted from the graph were used to construct a simplified version of the pangenome using haplotype 1 from *V. girdiana* as a reference (Fig. 6a). In this version of the graph, the representation of the sequence variations among the wild species was relative to *V. girdiana*. The reduction of the number of paths in the graph from 342 to 19 allowed us to prepare the pangenome for read mapping by significantly reducing the computational requirements. We indexed this reduced pangenome and mapped reads against it using vg. For each sample, >95% of the paired-end reads mapped onto the pangenome (Additional File 1: Table S18). After computing the snarls and the pack files from the graph, vg call was used to genotype the variants present in the graph (Fig. 6a).

**Fig. 6:**
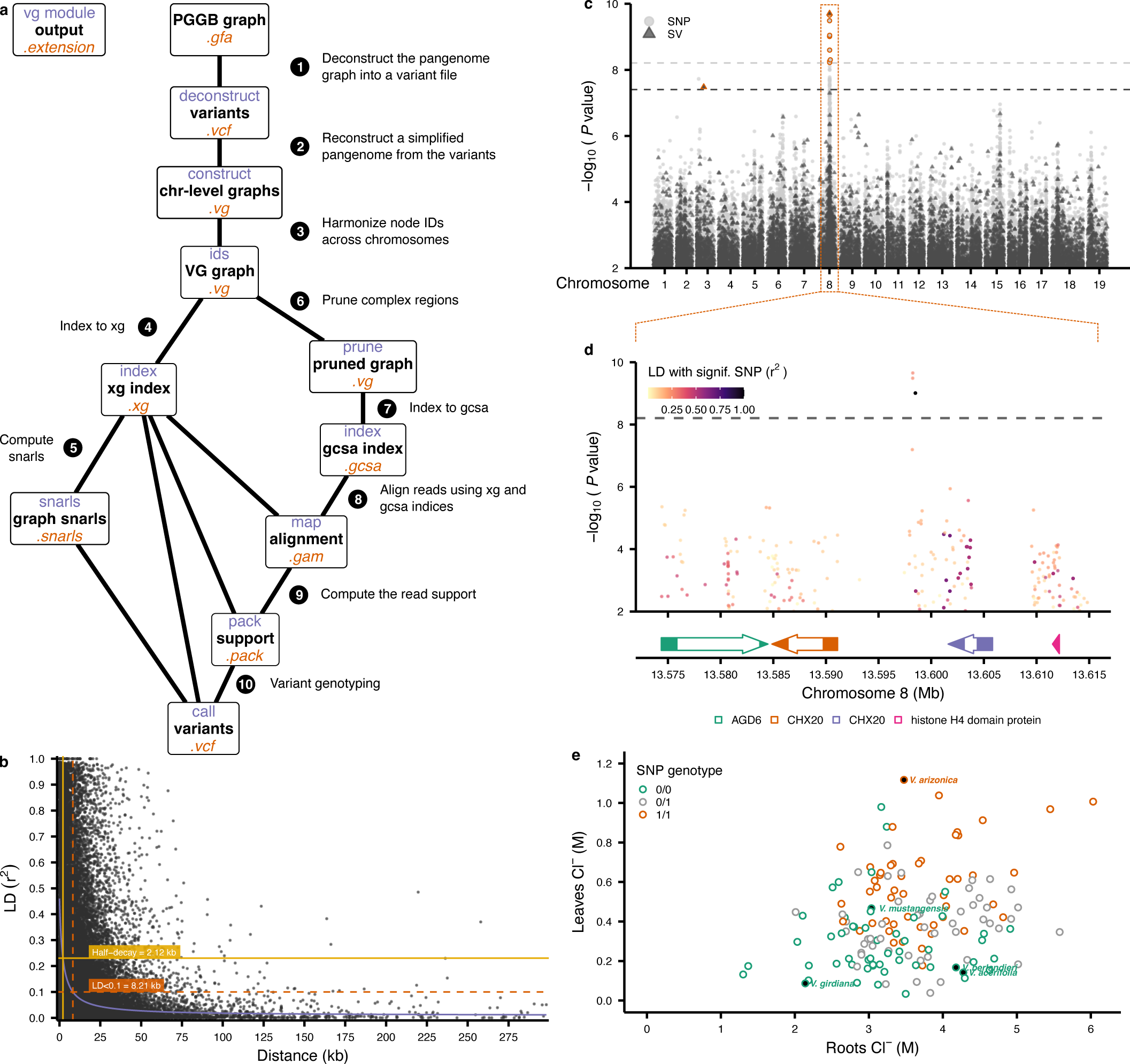
Pan-GWAS revealed significant associations with chloride concentration in the leaves. **a**, Workflow to generate the variant file required for the pan-GWAS. The name of the vg module used for each step is indicated in purple, the output type is in black, and the file extension is orange. The PGGB graph is deconstructed to a variant file using *V. girdiana* haplotype 1 as a reference (1). Vg construct is used to construct a pangenome from the VCF and the reference fasta (2). After merging the IDs (3), xg (4) and post-pruning (6) gcsa (7) indices are created to proceed with the alignments with vg map (8). The snarls are computed from the xg index (5) and the read support (pack) is extracted from the gam files (9) to finally proceed with the variant genotyping (10). **b**, Linkage disequilibrium decay in the SNP dataset used for the pan-GWAS. Half-decay is represented in yellow, while minimum decay with a threshold set to LD < 0.1 is colored in orange. The number of points was randomly down-sampled to 50k for visual representation. **c**, Manhattan plot representing significant association with the chloride concentration in the leaves. Significant associations are detected using a Bonferroni threshold set to −log10(0.05/n) and colored in orange. The SNPs are represented as light grey points, and the SVs as dark grey triangles. **d**, Manhattan plot focused on the significant region in chromosome 8 outlined by the orange box in **c** for the association with the leaf chloride concentration. The filling color is based on the pairwise linkage disequilibrium with the SNP located at chr08:13598495. The gene structural annotation of this region is represented at the bottom. **e**, Chloride concentrations (M) in the roots (x-axis) and in the leaves (y-axis) from the different samples used for the GWAS. Samples corresponding to the accessions assembled and used in the pangenome are filled in black. The colors represent the genotype at the significant SNP, homozygous reference (0/0) in green, heterozygous (0/1) in grey, and homozygous alternative (1/1) in orange.

After filtering, we examined the distribution of SNPs in the 19 paths. The SNPs exhibited rapid decay of linkage disequilibrium (LD), reaching half the maximum average r2 at 2.12 kb (Fig. 6b), consistent with previous reports in grapes [18,27]. We also performed pan-GWAS separately using 8,091,983 SNPs and 1,261,953 SVs, which should provide sufficient statistical power [28]. For both SNPs and SVs, a significant peak of association was detected on chromosome 8, and a single 1-bp deletion was found on chromosome 3 (Fig. 6c). Those genotyping results were supported by variant calling from a standard GWAS approach based on the *V. girdiana* reference (Additional File 2: Figure S9a).

The significant SNP detected in both pan-and standard GWAS was located in an intergenic region at chr08:13598495 (Fig. 6d) and was tightly associated with chloride concentration across samples (Fig. 6e). The SNPs in strong linkage disequilibrium with the significantly salt exclusion-associated SNP were located downstream, overlapping a gene locus spanning chr08:13601610- 13605816 (Fig. 6d). Based on a snpEff analysis, these SNPs appeared to be mostly missense variants, causing codons that produce different amino acids. Interestingly, the predicted protein from this region (VITVgdSC2_v1.0.hap1.chr08.ver1.0.g144890.t01) is a homolog of the *Arabidopsis thaliana* cation/H^+^ exchanger AtCHX20 (AT3G53720). The putative function of this gene was further supported by the presence of two notable InterPro domains: the Cation/H+ exchanger (IPR006153) and the Sodium/solute symporter superfamily (IPR038770). The gene upstream of the significant SNP was also found to be homolog to *AtCHX20* (Fig. 6d). Altogether, these results show that our super-pangenome allows the detection of base-level associations. We note, however, that when we investigated another related trait, root-chloride concentration, no significant association was detected with pan-GWAS (or, for that matter, with standard GWAS) (Additional File 2: Figure S9b,c).

## Discussion

Wild *Vitis* species represent a valuable genetic pool for grape cultivars and rootstock improvement. Despite an increasing number of *Vitis* genomes being published, no integrative analysis has been performed yet, limiting the exploration of such a diverse repertoire. Here we assembled the genomes of nine North American wild *Vitis* species to represent the diversity of the genus (Fig. 1), scaffolded them at chromosome level, and then phased haplotype (Table 1) to comprehensively assess the structural variations present in complex and repetitive regions in diploid genomes [29]. These accessions included species that contain characteristic agronomic traits, such as disease resistance and drought tolerance. The accessions also include *V. berlandieri*, *V. riparia,* and *V. rupestris*, the wild species trio responsible for the production of most of the grape rootstocks [30].

In recent years, pangenomes have been generated for numerous crop species. However, the concept of super-pangenomes is still in its early stages of development, although notable examples can be found in tomato [31], rice [32], sorghum [33], and potato [34]. Our super-pangenome differs from these previous efforts by using a reference-unbiased, graph-based approach. Its construction relied on all-vs-all sequence alignments of chromosome-level diploid assemblies thereby providing access with a nucleotide resolution to intra- and inter-specific genetic variants, beyond the limited gene space and overcoming reference-bias (Fig. 2). Our efforts provide an exemplar for future plant super-pangenomes and describe a pipeline that can be used in other species from the *de novo* assembly of diploid genomes to their pangenome construction. Our work paves the way for future research in *Vitis*. For example, this super-pangenome will be useful for additional comparative pangenomics analyses, and it will provide a backbone for including the wine-producing cultivars of *V. vinifera,* helping to identify and study variants of potential agronomic utility. More accessions could also be added for the species already present in this super-pangenome to develop species-level pangenomes. We selected one accession per species to build the super-pangenome since intra-species variation was beyond the scope of this work.

Our super-pangenome has provided insights into repetitive and genic content throughout the genus. For example, our characterization of the repetitive elements has shown that LTR Gypsy retrotransposons are enriched within private genomes (Fig. 2e). They are thus a predominant driving force behind the variable portion of the pangenome. TEs are known to impact the gene space in multiple ways. By their insertion near genes, they can regulate gene expression (Fig. 4h) and participate in epigenetic gene regulation [35]. TEs can also influence the gene content itself (Fig. 4i) by numerous mechanisms such as gene mutation, gene movement, but also duplication via unequal crossing over [35,36], which could all significantly contribute to the control of the variable genome size of a genus. Interestingly, the accurate representation of the sex-determining region in the pangenome graph evidenced an allele-specific pattern for the repeat distribution as previously reported in grapes [37]. The allele-specific nodes confirmed an accumulation of repetitive elements in the male allele, mechanism involved in the evolution of heteromorphic sex chromosomes [38]. Our results confirmed the significant contribution of repetitive elements notably TEs in the evolution of grape genomes and open the doors to further characterizations of the genus repetitive content. With respect to genes, we have found (unsurprisingly) that most genes are in the core genome. These genes are enriched for essential functions. In contrast to the core genome, the variable genome was significantly more impacted by TEs (Fig. 4) and SVs (Fig. 5) and contained a higher proportion of expanded/contracted gene families, potentially comprising the adaptative gene repertoire from the wild species. Including genomes from domesticated cultivars is likely to further expand the size of the dispensable genome, for two reasons. First, the dispensable genome increases with the addition of more samples (Fig 2a, 3a), and this trend is likely to continue based on the slopes predicted from our modeling until reaching a plateau for the gene content and most likely further increasing for the sequences as suggested for eukaryotic genomes [39]. Second, domesticated cultivars, which are clonally propagated, have much higher rates of genic hemizygosity (typically 10 to 15%) [14,40] compared to the < 5% found in the outcrossing species in this study (except for *V. arizonica*). Thus, cultivars may have a larger proportion of their genes captured in the dispensable genome.

We have demonstrated the utility and accuracy of the super-pangenome in three ways. First, by investigating hybrid genomes, we have shown that the super-pangenome accurately identified their parental origins and accurately characterized origin at the haplotype level. These results further support the broad potential and the flexibility of our pangenomics pipeline applicable in other species such as fruits crops (apples, pears, citrus, stone fruits trees, etc) by providing means to study the genetic variations between wild perennial accessions and the commercial hybrids derived from them. Second, we have recapitulated valuable genetic variants at the nucleotide level, specifically variants in the sex-determining region (Fig. 5f, Additional File 2: Figure S7) and that are associated with Pierce’s Disease (PD) (Additional File 2: Figure S8). The first led to the prediction of the flower sex phenotype of each accession by the presence/absence of nodes from the graph in the *VviINP1* gene, bypassing the need to perform manual gene annotation. For the latter, an interesting feature of the PD-associated variants is that they were more numerous in the variable (dispensable or private) genome compared to the core genome. This observation supports the notion that the variable genome contributes to species-specific adaptations, particularly those associated with biotic and abiotic stress, and it also consistent with the previous conjecture that the genetic basis for resistance may vary among *Vitis* species [25]. In the future, a pan-GWAS of PD-resistance across the genus may help reveal additional variants that contribute to this important trait.

Finally, we have demonstrated the utility of the super-pangenome by performing a pan-GWAS on chloride exclusion, leading to the identification of variants near gene loci homologs of *AtCHX20* on chromosome 8 (Fig. 6c,d). Interestingly, the SNPs were predicted to affect the amino acid composition of VITVgdSC2_v1.0.hap1.chr08.ver1.0.g144890 which could potentially have a significant impact on gene expression and protein activity [41]. In *V. vinifera* PN40024, the closest homolog is *Vitvi08g01174*, which is known to be induced by salt treatment [42]. In soybean (*Glycine max*), the closest homolog of VITVgdSC2_v1.0.hap1.chr08.ver1.0.g144890.t01 is GmNcl1 (Glyma.03g171500), which is also known to be induced by salinity [43]. Interestingly, the closest soybean homolog to AtCHX20 is GmSALT3 (Glyma.03g32900), the major salt tolerance gene in *G. max* [44]. Altogether, these results highlight the potential involvement of the *AtCHX20* homologs for salt tolerance in grape but would require further characterization to validate their functions. Thus, the pangenome has provided insights into a trait, salt tolerance, that is not only agronomically important but that is growing importance in the context of climate change and current agricultural uses. The current graph can be further augmented to embed the novel variants included within resequencing data, and it can be used as a reference for pan-GWAS based on germplasms that encompass multiple interfertile species. We, therefore, anticipate that the pangenome will be the backbone for many future applications, and particularly for unraveling the genetic basis of critical agronomic traits across *Vitis*.

## Conclusions

In summary, we constructed a super-pangenome to represent and analyze North American wild species of the *Vitis* genus. The assembly of phased diploid genomes for the nine selected species was a fundamental starting point to ensure an accurate representation of genetic variations that occur within and between genomes. We were able to navigate through different layers of information integrated within the pangenome graph from large structural variations to single nucleotide polymorphisms. We investigated and interpreted the genomic diversity within the genus and provide novel insights regarding the core and the variable genome in grapes. We expect that many more insights on *Vitis* evolution as the pangenome for this genus grows and integrates more species, notably from distinct geographical regions such as Europe and Asia, which should help to further elucidate grapevine domestication for example. The integration of the *V. vinifera* species will provide direct insights into the genetic bases of the organoleptic properties of the wine-producing berries [45]. The concept of pangenomics provides many opportunities to finally move from a single-reference to a reference-free approach that incorporates broad genetic diversity when comparing different genomes. Significant improvements in accuracy are expected for approaches sensitive to single nucleotide changes such as transcriptomics and genetic association studies of populations with frequent inter-species introgressions.

## Methods

### Plant material and tissue collection

Several plant tissues (see corresponding sections below) were collected for sequencing of DNA, RNA, and full-length transcripts, Bionano next-generation mapping, and flow cytometry. All plant material intended for nucleic acid extraction was immediately frozen upon collection and ground in liquid nitrogen. Most of the samples were collected from accessions maintained in the vineyards by the Foundation Plant Services at the University of California, Davis (Additional File 1: Table S1). The selection of the different accessions was based on plant material availability and agronomical interests evidenced previously.

### Library preparation and sequencing

High-molecular-weight genomic DNA (gDNA) was isolated from young leaves using the method described in Chin et al. 2016 [46]. DNA purity was evaluated with a Nanodrop 2000 spectrophotometer (Thermo Scientific, IL, USA), DNA quantity with the DNA High Sensitivity kit on a Qubit 2.0 Fluorometer (Life Technologies, CA, USA), and DNA integrity by pulsed-field gel electrophoresis. gDNA was cleaned with 0.45x AMPure PB beads (Pacific Biosciences, CA, USA) before library preparation. SMRTbell template was prepared with 15 µg of sheared DNA using SMRTbell Express Template Prep Kit (Pacific Biosciences, CA, USA) following the manufacturer’s instructions. SMRTbell template was size selected using the Blue Pippin instrument (Sage Science, MA, USA) using a cutoff size of 25-80 kb. The size-selected library was cleaned with 1x AMPure PB beads. The SMRTbell library was sequenced on a PacBio Sequel II platform using V2.0 chemistry (DNA Technology Core Facility, University of California, Davis, CA, USA) (Additional File 1: Table S2).

Ultra-high molecular weight DNA (> 500 kb) was also extracted from young fresh leaves using Amplicon Express (Pullman, WA, USA). After labeling with a DLE-1 non-nicking enzyme (CTTAAG) and staining following the instructions of the Bionano PrepTM Direct Label and Stain Kit (Bionano Genomics, CA, USA), DNA was loaded onto a SaphyrChip nanochannel array for imaging with the Saphyr system (Bionano Genomics, CA, USA) (Additional File 1: Table S3).

For *V. acerifolia*, *V. arizonica*, *V. monticola,* and *V. riparia*, DNA-Seq libraries were prepared from 1 mg of DNA extracted from young leaves using the Kapa LTP library prep kit (Kapa Biosystems, MA, USA). After quantity and quality evaluation with the High Sensitivity chip of a Bioanalyzer 2100 (Agilent Technologies, CA, USA), libraries were sequenced in 150-bp-long paired-end reads on an Illumina HiSeq 4000 (DNA Technology Core Facility, University of California, Davis, CA, USA) (Additional File 1: Table S5). For *V. rupestris*, the reads were produced using a HiSeq X Ten instrument (IDSeq, CA, USA). For the four other genotypes, the sequencing data were retrieved from the BioProject PRJNA731597 [4].

Total RNA was extracted from young leaves as described in Cochetel et al. (2021) [47] using the Cetyltrimethyl Ammonium Bromide (CTAB)-based protocol from Blanco-Ulate et al. (2013) [48]. RNA purity was evaluated using a Nanodrop 2000 spectrophotometer (Thermo Scientific, IL, USA). DNA quantity was verified using a Qubit 2.0 Fluorometer and a broad range RNA kit (Life Technologies, CA, USA). Finally, the RNA integrity was evaluated through electrophoresis and with an Agilent 2100 Bioanalyzer (Agilent Technologies, CA, USA). Two types of cDNA libraries were produced. For short reads, the Illumina TruSeq RNA sample preparation kit v.2 (Illumina, CA, USA) was used for library preparation following the low-throughput protocol. Library quantity and quality were evaluated with the High Sensitivity chip in an Agilent Bioanalyzer 2100 (Agilent Technologies, CA, USA). The sequencing was performed using an Illumina HiSeq 4000 (DNA Technology Core Facility, University of California, Davis, CA, USA) to produce 100-bp-long paired-end reads. For full-length cDNA sequencing, cDNA SMRTbell libraries were prepared. The first-strand synthesis and cDNA amplification were performed with the NEB Next Single Cell/Low Input cDNA Synthesis & Amplification Module (New England, MA, USA). The cDNA was then purified with ProNex magnetic beads (Promega, WI, USA) according to the Iso-Seq Express Template Preparation for Sequel and Sequel II Systems protocol (Pacific Biosciences, CA, USA) (Additional File 1: Table S4). Long cDNA (> 2kb) fragments were selected with ProNex magnetic beads (Promega, WI, USA), and at least 80 ng were used to prepare the cDNA SMRTbell library. DNA damage repair and SMRTbell ligation were performed using the SMRTbell Express Template Prep kit 2.0 (Pacific Biosciences, CA, USA). One SMRT cell per genome was sequenced on the Sequel I platform (DNA Technology Core Facility, University of California, Davis, CA, USA) (Additional File 1: Table S4).

### Genome assembly, phasing, and chromosome-scaling

For each genome, SMRT sequences (Pacific Biosciences, CA, USA) were assembled into haplotype-resolved contigs using the diploid-aware assembler Falcon-Unzip [46], as described in Massonnet et al., 2020 [6] (Additional File 1: Table S2), and polished using Arrow from ConsensusCore2 v.3.0.0 (https://github.com/PacificBiosciences/ccs). Long imaged molecules (>150 kb) from the next-generation mapping (Bionano Genomics, CA, USA) were assembled using Bionano Solve v.3.3 [49] with the parameters described in Cochetel et al., 2021 [47]. Using HybridScaffold v.04122018 [49] with the conflict resolution parameters “-B2 −N1”, hybrid assemblies were generated through the scaffolding of the Arrow-polished contigs with the consensus genome maps. Despite four next-generation mappings for *V. monticola*, we were not able to generate a consensus genome map for this genome. For *V. riparia*, no optical maps were available at the time of the assembly (Additional File 1: Table S3). Scaffolds were simultaneously phased and separated into chromosomes using the tool suite HaploSync [12] and the ∼2,000 rhAmpSeq *Vitis* markers [13] (Additional File 1: Table S19). The genome of *V. arizonica* was already released to support another study [25]. Illumina DNA-Seq reads were used during the chromosome reconstruction for coverage analysis (Additional File 1: Table S5). Gene space completeness was evaluated using BUSCO v.3.0.2 scores [50]. In addition, CDS from PN40024 [5] were mapped on each genome using pblat v.36×2 [51] with the parameters “maxIntron=35000-minIdentity=0”. Before the alignments, the set of PN40024 CDS used was filtered to keep only the sequences uniquely mapping on PN40024 itself with a threshold fixed at 80% for identity and coverage. After the alignments on the nine species, hits were filtered with 50% identity and coverage, but also a minimum overlap of 80% on the reference gene loci (Additional File 1: Table S9). To evaluate heterozygosity, NUCmer from the tool MUMmer v.4.0.0beta5 [23] was used with the parameter “--mum” to align the haplotypes of each genome against each other. All the resulting variant types and lengths were concatenated together to estimate the proportion of the genome covered by variants (Additional File 1: Table S7). For representation per variant type (Additional File 2: Figure S1), translocations, inversions and any variants not classified as SNP or INDEL were merged into “other” category. Inter-species variations were also computed aligning the haplotypes of the different species against each other and using the same metrics as for the heterozygosity estimation (Additional File 1: Table S11).

### Structural and functional annotation

The structural annotation of the protein-coding gene loci was performed using an extensive ab-initio prediction pipeline described in Cochetel et al. (2021) [47], available at https://github.com/andreaminio/AnnotationPipeline-EVM_based-DClab. Repeats were annotated using RepeatMasker v.open-4.0.6 [52]. For each assembly, the proportion of bases covered by repetitive elements were represented as repeat percentages. RNA-Seq reads were filtered based on quality and adapters were trimmed using Trimmomatic v.0.36 [53]. Quality controls pre- and post-trimming were performed with fastQC [54]. Filtered reads were used to produce transcriptome assemblies through Stringtie v.1.3.4d [55] and Trinity v.2.6.5 [56]. For the IsoSeq reads, IsoSeq v.3 (https://github.com/PacificBiosciences/IsoSeq) was used for the extraction, and LSC v.2.0 [57] for the polishing of the low-quality isoforms. For *V. rupestris*, IsoSeq reads were not available at the time of the annotation, RNA-Seq was used as transcriptomic evidence (Additional File 1: Table S4). High-quality gene models were then generated using PASA v.2.3.3 [58] from the transcript evidence and external databases. The same input was used to produce genome alignments with Exonerate v.2.2.0 [59], PASA, and MagicBLAST v.1.4.0 [60]. Ab initio predictions were also generated using Augustus v.3.0.3 [61], BUSCO v.3.0.2 [50], GeneMark v.3.47 [62], and SNAP v.2006-07-28 [63]. Based on all the above evidence, EvidenceModeler v.1.1.1 [64] was used to generate consensus gene models. The functional annotation was produced with Blast2GO v.4.1.9 [65] combining blastp v.2.2.28 [66] or diamond v.2.0.8 [67] hits against the Refseq plant protein database (https://ftp.ncbi.nlm.nih.gov/refseq/, retrieved January 17th, 2019) and InterProScan v.5.28-67.0 [68] results. Hemizygosity was estimated through the alignments of the CDS of each haplotype per genome against each other using GMAP v.2019.09.12 [69]. Results were filtered with a threshold of 80% for identity and coverage. To prevent assembly bias, CDS were also aligned on the unplaced sequences (Additional File 1: Table S8). Hmmsearch from HMMER v.3.3 (http://hmmer. org/) was used to scan the proteomes using NLR-specific Pfam domains [70]; PF00931, PF01582, PF00560, PF07723, PF07725, and PF12799. For proteins containing coiled-coil (CC) domains, annotations were extracted from the InterProScan results. NLR-annotator [71] was used as an additional source of evidence to identify NLR genes. After identifying NBS-LRR genes missing the NB-ARC domain (Pfam PF00931) as TIR-X or CC-X, eight classes were defined with CC-NBS-LRR, CC-NBS, TIR-NBS-LRR, TIR-NBS, NBS-LRR, and NBS. For gene ontology enrichment analysis, the R package topGO v2.48.0 [72] was used with an FDR-adjusted *P* value threshold set to 0.01.

### Flow cytometry

DNA content was estimated using flow cytometry as described in Cochetel et al. (2021) [47]. The nuclei extraction was performed using the Cystain PI absolute P kit (Sysmex America Inc., IL, USA). For the internal reference standard, *Lycopersicon esculentum* cv. Stupické polní tyčkové rané was selected, with a known genome size of 2 C = 1.96 pg; 1 C = 958 Mbp [73]. Young leaves (∼ 5 mg) from grape and tomato were finely cut with a razor blade in a Petri dish containing 500 mL of extraction buffer and filtered through a 50 mm filter (CellTrics, Sysmex America Inc., IL, USA). Propidium iodide staining solution (2 mL) was then added to the nuclei suspension [74,75]. Measurements were acquired with a Becton Dickinson FACScan (Franklin Lakes, New Jersey) equipped with a 488 nm laser. Raw data were imported and analyzed with the R package flowPloidy v.1.22.0 [76] (Additional File 1: Table S9). For *V. berlandieri*, *V. riparia*, and *V. rupestris*, estimates from the same species were retrieved from literature [77].

### Pangenome graphs construction

A nucleotide-level super-pangenome graph was built using the tools from the PGGB pipeline (https://github.com/pangenome/pggb) [11,21]. To test different configurations of the aligner (wfmash) and the graph builder (seqwish), chromosome 1 of the first haplotype from *V. acerifolia* and *V. berlandieri* were selected arbitrarily (Additional File 1: Figure S9). For wfmash, the segment length (1, 5, 10, 20, 50, 100kb) and the identity (70, 75, 80, 85, 90, 95%) were evaluated and alignments were converted to a graph using a fixed kmer size = 19 bp for seqwish. After the final selection of wfmash parameters “−s 10000 −p 85”, the kmer size was tested from 15 to 405 bp and the default 49 bp was selected. The homologous chromosome sequences of each haplotype were aligned against each other between (e.g. chromosome 1 of species A haplotypes aligned against chromosome 1 of species B haplotypes) and within (e.g. chromosome 1 of species A haplotype 1 aligned against chromosome 1 of species A haplotype 2) species using wfmash (https://github.com/waveygang/wfmash) with the parameters “−p 85 −s 10000 −n 1”. The unanchored sequences from each genome were not included. The alignments were merged per chromosome and processed with seqwish v0.7.3 [21] to produce the pangenome graphs with “-k 49”. Finally, two passes of smoothxg (https://github.com/pangenome/smoothxg) were performed to polish the pangenome. The parameters for the first pass were the following: “-w 68017 −K −X 100 −I 0.85 −R 0 −j 0 −e 0 −l 4001 −P “1,4,6,2,26,1” −O 0.03 −Y 1700 −d 0 −D 0 −V” and the parameters for the second were: “-w 76619 −K −X 100 −I 0.85 −R 0 −j 0 −e 0 −l 4507 −P “1,4,6,2,26,1” −O 0.03 −Y 1700 −d 0 −D 0 −V”. These options were derived from the all-in-one PGGB command (https://github.com/pangenome/pggb) using target poa lengths (-l) of 4001 and 4507 for the first and second smoothxg pass, respectively. The chromosome-level graphs were joined using the module “ids” from vg v.1.48.0 [22] to produce a non-overlapping node id space. In the resulting pangenome, each chromosome from the 18 genomes was represented as a path. For each path, node ids and sequences were extracted and categorized in the core, dispensable, or private genomes based on their presence among species. The source code used to build and analyze the pangenome was released on GitHub [78] and referenced on Zenodo [79]. A second pangenome was built adding to the later the genomes of three hybrid species; 101-14 Millardet et de Grasset (101-14 Mgt: *V. riparia* x *V. rupestris*), Richter 110 (110R: *V. berlandieri* x *V. rupestris*), and Kober 5BB (*V. berlandieri* x *V. riparia*)[7]. Wfmash and seqwish parameters were the same. For smoothxg, the parameters were modified to take into account the high number of genomes; for the first pass “-w 92023 −K −X 100 −I 0.85 −R 0 −j 0 −e 0 −l 4001 −P “1,4,6,2,26,1” −O 0.03 −Y 2300 −d 0 −D 0 −V” and the second pass “−w 103661 −K −X 100 −I 0.85 −R 0 −j 0 −e 0 −l 4507 −P “1,4,6,2,26,1” −O 0.03 −Y 2300 −d 0 −D 0 −V”.

### Orthology analysis

Transcripts from the nine species were aligned on the genome of each other using pblat v.36×2 [51] with the parameters “maxIntron=35000-minIdentity=0”. After filtering with a threshold of 80% for identity and coverage, hits were summarized at the gene level. To evaluate colinearity between genomes, the above pblat hits were further filtered to cover at least 80% of the reference gene loci. Colinear blocks were then detected using McScanX_h from McScanX v.11.Nov.2013 [80] with default parameters.

### Transcript abundance quantification

Salmon v.1.10.1 [81] was used for transcript-level quantification. To get high-accuracy alignments, decoys were generated using the genome sequences in addition to the CDS (https://combine-lab.github.io/alevin-tutorial/2019/selective-alignment/). Decoy regions from the genome are used during the scoring of the alignments to reduce false mapping of sequenced fragments that originate from unannotated genomic loci [82]. The indexes were created using a kmer size of 31 bp. Transcript levels were then quantified with the parameters “--seqBias –gcBias--validateMappings” with the paired-end RNA-Seq data used for the gene annotation. Quantification results were imported and TPM values were extracted using the R package tximport v.1.24.0 [83].

### Population structure

DNA-Seq samples were sequenced as described previously [4], raw sequencing reads were deposited on NCBI under the BioProject PRJNA984685 [84], and previously sequenced reads [4,25] were retrieved from the BioProjects PRJNA731597 [85] and PRJNA842753 [86]. DNA-Seq reads were quality-based trimmed using trimmomatic and mapped against the haplotype 1 of an equidistant reference genome from the *Vitis vinifera* clade, Cabernet Sauvignon [6], using bwa v.0.7.17-r1188 [87] and the parameter “mem”. Variant calling per sample was performed using the bcftools v1.17 [88] commands mpileup and call with the parameters “--keep-alts--gvcf 0 -- multiallelic-caller”. The VCF files were merged using bcftools merge “--gvcf”. Variant sites and samples were filtered using bcftools filter “-e ‘F_MISSING > 0.25 || MAF <= 0.01’”, discarding 19 samples (153 remaining). Samples were further filtered to be representative of their groups and avoid any over-representation of a species/group overall. The maximum number of representative species per group was set to 10 with a selection based on coverage. The inclusion was ensured for the samples of the assembled genomes, a filter on coverage >= 10x was used, and this led to the final selection of 90 samples. Bedtools v.2.30.0 [89] intersect was used to discard variant sites in repetitive regions, further filtered with bcftools filter “-e ‘AC==0 || AC==AN’ --SnpGap 10”, bcftools view “-m2 −M2 −v snps”, bcftools filter “−i ‘QUAL>=30’”, bcftools view “−e ‘QUAL < 30 || MQBZ < −3 || RPBZ < −3 || RPBZ > 3 || SCBZ > 3 || INFO/DP<5 || INFO/DP>2000’” and pruned with plink v.1.90b6.21 [90] “--indep-pairwise 50 5 0.2”. The population stratification was finally evaluated by dimension reduction using plink “--pca”.

### Phylogenetic analysis

Proteomes from the representative haplotype 1 of each genome were compared to each other using OrthoFinder v.2.5.4 [91]. Data for *Muscadinia rotundifolia* cv. Trayshed [47] were retrieved from http://www.grapegenomics.com/. The sequences of each single-copy gene orthogroups were aligned using MUSCLE v.5.1 [92]. Alignments were concatenated and parsed with Gblocks v.0.91b [93] including positions with gap allowed in < 50% of the sequences. The evolutionary model was selected using the results from ModelTest-NG v.0.1.7 [94] The phylogenetic tree was generated using RAxML-NG v.1.1 [95] (Additional File 2: Figure S1). The Maximum Likelihood (ML) method was selected with the optimized evolutionary model JTT+I+G4+F, using ten parsimony starting trees, and a bootstrapping of 1,000 replicates. For clock calibration, the single copy orthologs were used as input for BEAUti v.1.10.4 [96] to generate the preliminary XML file for BEAST v1.10.4 [96] analysis. A monophyletic partition was set for the North American species in the subgenus *Vitis*. A calibration point was established at the crown age of this subgenus which was estimated to be 17.8 million years ago [97], with a normal distribution and standard deviation of 1. Ten independent Markov chain Monte Carlo (MCMC) chains, each consisting of 1,000,000 generations, were run using BEAST. The JTT substitution model, with four Gamma categories, a strict clock, and the Birth-Death Model with a random starting tree were used. Sampling was performed every 1,000 generations. The resulting log and trees files were combined using LogCombiner v.1.10.4 [96], and the maximum clade credibility tree was generated using TreeAnnotator v.1.10.4 [96] with a burn-in of 10,000 generations (Fig. 1k). For the dN/dS estimations, proteins from each haplotype used to construct the pangenome were separated into core and dispensable proteins selecting the longest isoform as representative. The orthogroups identification and their sequence alignments were performed as described above for the whole proteomes and converted to CDS using PAL2NAL v.13 [98]. For the core proteins, the tree computed earlier with the representative haplotype 1 was used. For the dispensable proteins, a tree was computed for each orthogroup using RAxML-NG with the same parameters as for the proteomes. Using the CDS alignments and the phylogenetic trees, the dN/dS ratio was estimated for every core and dispensable orthogroups using codeml from PAML v.4.10.0 [99] following the best practice guidelines [100]. For every gene of haplotype 1, a representative protein isoform was selected based on pfam domain coverage to study gene family evolution. They were aligned between genomes using all-vs-all diamond blastp with parameters “--sensitive --evalue 0.000001”. The gene family clusters were defined using Markov clustering (mcl) [101] with an inflation value set to 3 resulting in the identification of 9,220 gene families. Using the previously clock-calibrated phylogenetic tree, a computational analysis of gene family evolution was performed using CAFE v.5.0 [102] using default settings and a p-value threshold of 0.01. The lambda parameter specific to this dataset was 0.0103. After filtering for significant expansion or contraction, 450 gene families were retained.

### Structural variant analysis

Single-nucleotide polymorphisms and structural variants were extracted from the pangenome graph using vg deconstruct and filtered to only consider top-level variants (LV=0). Multi-allelic sites present in the VCFs were normalized to bi-allelic sites using “norm” from bcftools with the parameter “-m-any”. Files were divided into SNPs and INDELs using bcftools view and the parameter “--type”. While the variant types MNP (Multiple Nucleotide Polymorphism) and other were also detected in the vcf file, they were not considered further as they correspond to complex variation scenarios from the decomposition of the graph into variants. For the comparison of haplotypes in each genome, the vcf were filtered to only retain variants present in the alternative haplotype for each haplotype considered as reference. Insertions and deletions were extracted from the INDELs files using bcftools filter with “--include ‘strlen(REF)<strlen(ALT)’” and “--include ‘strlen(REF)>strlen(ALT)’”, respectively. SnpEff v.5.1 [24] was used to annotate the functional impact of the different types of variants. The unfiltered VCFs were compared with the inter-genomic pairwise comparisons made with NUCmer. For every reference genome, the insertion sites and deletions were concatenated into INDELs. The SNPs and INDELs detected from the different query genomes were combined per reference genome into a non-overlapping bed file using bcftools merge summarizing the entire proportion of the reference genome impacted by variants. The variant sites identified from vg deconstruct were compared with these SNPs and INDELs reference datasets using bedtools intersect. An enrichment analysis was performed on the different variant effects by comparing the proportion of variants with effect between the genes considered hemizygous and the total set of genes impacted by variants. A two-sided Fisher test was performed followed by a Bonferroni correction of the p-values. Effects presenting an enrichment > 1 and a p-adjusted value < 0.05 were considered.

### Sex-determining region analysis

The sex-determining region (SDR) was localized in all the genomes using GMAP. Using *V. arizonica* as a reference, as its SDR was manually curated for a previous study [6], we discarded haplotypes that would require manual curation of their assembly and kept 11 female and 4 male haplotypes. We selected a sub-region from the gene encoding VviYABBY3 to the WRKY transcription factor to focus between boundaries that contain most of the female/male alleles differences [6]. For each haplotype, variant sites generated from the graph decomposition (vg deconstruct) were selected if they are present in all the haplotypes of the alternative allele considered. Allele specific nodes were defined as nodes present in all the haplotype of an allele and absent in the haplotypes of the alternative. Repeats overlapping any allele-specific nodes were considered. For the gene representation, the annotation of *V. arizonica* was used.

### Pan- and classic GWAS

The unfiltered VCFs from vg deconstruct were used with the reference genome fasta of the haplotype 1 of *V. girdiana* SC2 as inputs to the command “construct” from vg. One graph was constructed per chromosome and the id space of the 19 graphs was merged using vg ids. The gcsa index and the xg index were computed with vg index to proceed with the alignments using vg map. The read support was then extracted from the alignment files using vg pack. Finally, the variant genotyping was performed using the following command structure per sample: “vg call –threads number_of_cores −k sample.on.graph.pack −r graph.snarls −s sample −a graph.xg > sample.on.graph.vcf”. The flag −a is essential to merge all the samples’ VCFs as described in https://github.com/vgteam/vg#calling-variants-using-read-support. The 153 samples were merged into a single VCF using the command “merge –merge all” from bcftools. An initial filter was applied to discard variant sites from the VCF if they were presenting more than 25% of missing genotypes or a minor allele frequency lower than or equal to 1% using bcftools filter. SNPs sites were extracted using bcftools view and the parameter “-v snps” while the SVs were obtained using bcftools view “–exclude-types snps”. The different chromosome-level VCF files were concatenated together using bcftools concat. After quality assessment, VCF files were further filtered with bcftools filter “-e ‘MAF <= 0.05 || QUAL < 30 || DP < 153 || DP > 5000’”. Each variant site was renamed with bcftools annotate using the parameter “–set-id +’%CHROM\_%POS’”. For linkage disequilibrium (LD) analysis, the following command was used to prune the dataset towards a subset of markers in approximate linkage equilibrium using plink: “plink –vcf name.vcf –double-id –allow-extra-chr −r2 gz –maf 0.1 –geno 0 –mind 0.1–ld-window 10 –ld-window-kb 300 –ld-window-r2 0 –out name –thin-count 500000”. LD decay was evaluated using the model from Hill and Weir (1988) [103] as previously described in grapes [104]. For GWAS, plink was used to prune the files using “–indep-pairwise 50 5 0.2” required to generate the standardized relatedness matrix to account for population structure with gemma v.0.98.3 [105]. The GWAS was performed using the matrix and the unpruned variant datasets with linear mixed models (LMMs) to test for association using gemma.

A classic GWAS was also performed using the haplotype 1 of *V. girdiana* SC2 as a reference. The DNA-Seq reads were mapped using bwa and the parameter “mem”. The alignments were prepared for GATK using several commands from the Picard toolkit (https://github.com/broadinstitute/picard) starting with the validation of the files using “ValidateSamFile”. A sequence dictionary for the reference genome was obtained using “CreateSequenceDictionary”. The commands “MarkDuplicates”, “AddOrReplaceReadGroups” and “BuildBamIndex” were used to complete the file preparation. The variant calling was performed using GATK v.4.0.12.0 [106] with the following command: HaplotypeCaller −I aln.bam –base-quality-score-threshold 20 –sample-ploidy 2 –native-pair-hmm-threads 4 −R ref.fasta −ERC GVCF –disable-read-filter GoodCigarReadFilter –output aln.g.vcf −L chr_name”. The single sample gVCFs were imported into a GenomicsDB before the joint genotyping using “GenomicsDBImport” and the final VCF was generated using “GenotypeGVCFs”. The VCF was filtered following GATK’s guidelines using “bcftools filter −e ‘QD < 2 || FS > 60 || SOR > 3 || MQ < 40 || MQRankSum < −12.5 || ReadPosRankSum < −8’” (https://gatk.broadinstitute.org/hc/en-us/articles/360035890471-Hard-filtering-germline-short-variants). The separation into SNPs and SVs, the last filterings and the GWAS followed the same steps as for the pan-GWAS.

### Data visualization and statistical analysis

Rstudio v.2023.03.1.446 [107] running R v.4.2.1 [108] was used to run the R packages cited above and all the figures were generated mostly using the package tidyverse v.2.0.0 [109]. Genomics data were parsed using GenomicFeatures v.1.48.4 [110], phylogenetic trees were drawn using ggtree v.3.4.4 [111] after importing the data with treeio v.1.20.2 [112], the upset plot was generated using UpSetR v.1.4.0 [113], the gene representations were obtained using gggenes v.0.5.0 [114], and the chord diagrams were generated using circlize v. 0.4.15 [115].

### Declarations

#### Ethics approval and consent to participate

Not applicable.

### Consent for publication

Not applicable.

### Availability of data and materials

All the raw sequencing data are available as an NCBI BioProject under the accession number PRJNA984685 [84]. Some of the DNA-seq data from the natural populations of wild grape species previously published [4,25] were accessed from the BioProjects PRJNA731597 [85] and PRJNA842753 [86]. The genome files associated with *V. arizonica* were already released in a previous work [25]. Genome assemblies, annotations, and the associated genome browser are accessible at http://www.grapegenomics.com/, files released in previous work [47] for *Muscadinia rotundifolia* cv. Trayshed were retrieved from the same source. Optical maps, genome files, and pangenome graphs were uploaded on Zenodo [116]. Code and scripts used in the construction and analysis of the pangenome are available on GitHub under MIT License [78] and on Zenodo [79].

## Authors’ contributions

D.C. and N.C. designed the study. R.F.-B. processed the samples, performed the nucleic acids extractions, and prepared the sequencing libraries. R.F.-B. and T.K. performed the flow cytometry measurements, N.C. and T.K. analyzed the results. A.M. assembled the genomes and annotated *V. rupestris*. N.C. scaffolded into chromosomes, phased and annotated all the genomes. N.C., A.G. and E.G. built the pangenome. N.C. and J.F.G. performed the phylogenetic analysis. N.C. performed all the other bioinformatic analyses under the supervision of D.C.. N.C. and D.C. wrote the original manuscript draft, the co-authors reviewed and edited the manuscript, N.C., D.C. and B.S.G. wrote the final version. All authors read and approved the final manuscript.

## Acknowledgments

The authors acknowledge Victor Llaca (Corteva Agriscience), Lance Cadle-Davidson (USDA-ARS), and Bruce Reisch (Cornell University) for the *V. rupestris* optical maps.

## Supplementary Information

**Additional File 1**: **Table S1**. Accessions collected for the nine Vitis species. **Table S2**. PacBio Single Molecule Real-Time sequencing statistics. **Table S3**. Consensus genomes maps statistics. **Table S4**. RNA sequencing statistics. **Table S5**. Short-read DNA sequencing statistics. **Table S6**. N50 (bp) statistics at chromosome level. **Table S7**. Heterozygosity estimates from nucmer structural variants analysis. **Table S8**. Hemizygosity estimates from pairwise gene mapping analysis using GMAP. **Table S9**. Genome size estimations from flow cytometry and ploidy estimations using PN40024 gene representation. **Table S10**. Gene ontology enrichment analysis in the rapidly evolving gene families. **Table S11**. Structural variants coverage from pairwise assembly comparisons with nucmer. **Table S12**. Absolute and relative number of genes per class including the ambiguous class. **Table S13**. Absolute and relative number of genes per class without the ambiguous class. **Table S14**. Consistency statistics between the graph-inferred and the orthology-based gene pangenomes. **Table S15**. Proportion of rapidly evolving genes per pangenome class. **Table S16**. Consistency statistics between the variants identified with nucmer (inter-genome pairwise comparisons) and with vg deconstruct (pangenome graph). **Table S17**. SNP frequency estimates from graph decomposition using vg deconstruct. **Table S18**. Mapping on the pangenome statistics (vg map). **Table S19**. HaploSync runs.

**Additional File 2: Figure S1.** Frequency of variants between the haplotypes of each genome after intra-genomic comparisons. **Figure S2**. Graph-based pangenome modeling. Figure S3. Transposable elements composition in the pangenome. **Figure S4.** Comparison of the final pangenome sizes. **Figure S5.** NBS-LRR gene distribution. **Figure S6**. Frequency of the variant type modifier in the different pangenome classes. **Figure S7**. Structural variants at the sex-determining region within the *VviINP1* gene. **Figure S8**. Consistency of the detected variants in the pangenome graph with the previously PDR-associated regions. **Figure S9**. Single-reference GWAS and root chloride association analysis. **Figure S10**. PGGB modules optimization.

## References

1. Alston JM, Sambucci O. Grapes in the World Economy. In: Cantu D, Walker MA, editors. The Grape Genome. Springer International Publishing; 2019. p. 1–24.

2. Rahemi A, Dodson Peterson JC, Lund KT. Grape Rootstocks and Related Species. Springer International Publishing; 2022.

3. Walker MA, Heinitz C, Riaz S, Uretsky J. Grape Taxonomy and Germplasm. In: Cantu D, Walker MA, editors. The Grape Genome. Springer International Publishing; 2019. p. 25–38.

4. Morales-Cruz A, Aguirre-Liguori JA, Zhou Y, Minio A, Riaz S, Walker AM, et al. Introgression among North American wild grapes (Vitis) fuels biotic and abiotic adaptation. Genome Biol. 2021;22:254.

5. Jaillon O, Aury JM, Noel B, Policriti A, Clepet C, Casagrande A, et al. The grapevine genome sequence suggests ancestral hexaploidization in major angiosperm phyla. Nature. 2007;449:463–8.

6. Massonnet M, Cochetel N, Minio A, Vondras AM, Lin J, Muyle A, et al. The genetic basis of sex determination in grapes. Nat Commun. 2020;11:2902.

7. Minio A, Cochetel N, Massonnet M, Figueroa-Balderas R, Cantu D. HiFi chromosome-scale diploid assemblies of the grape rootstocks 110R, Kober 5BB, and 101–14 Mgt. Sci Data. 2022;9:660.

8. Khan AW, Garg V, Roorkiwal M, Golicz AA, Edwards D, Varshney RK. Super-Pangenome by Integrating the Wild Side of a Species for Accelerated Crop Improvement. Trends in Plant Science. 2020;25:148–58.

9. Lei L, Goltsman E, Goodstein D, Wu GA, Rokhsar DS, Vogel JP. Plant Pan-Genomics Comes of Age. Annu Rev Plant Biol. 2021;72:411–35.

10. Wang S, Qian Y-Q, Zhao R-P, Chen L-L, Song J-M. Graph-based pan-genomes: increased opportunities in plant genomics. Usadel B, editor. Journal of Experimental Botany. 2023;74:24– 39.

11. Garrison E, Guarracino A, Heumos S, Villani F, Bao Z, Tattini L, et al. Building pangenome graphs. [cited 2023 May 5]; Available from: https://www.biorxiv.org/content/10.1101/2023.04.05.535718v1

12. Minio A, Cochetel N, Vondras AM, Massonnet M, Cantu D. Assembly of complete diploid-phased chromosomes from draft genome sequences. G3 Genes|Genomes|Genetics. 2022;12:jkac143.

13. Zou C, Karn A, Reisch B, Nguyen A, Sun Y, Bao Y, et al. Haplotyping the Vitis collinear core genome with rhAmpSeq improves marker transferability in a diverse genus. Nat Commun. 2020;11:413.

14. Zhou Y, Minio A, Massonnet M, Solares E, Lv Y, Beridze T, et al. The population genetics of structural variants in grapevine domestication. Nat Plants. 2019;5:965–79.

15. Velasco R, Zharkikh A, Troggio M, Cartwright DA, Cestaro A, Pruss D, et al. A High Quality Draft Consensus Sequence of the Genome of a Heterozygous Grapevine Variety. Dilkes B, editor. PLoS ONE. 2007;2:e1326.

16. Girollet N, Rubio B, Bert P-F. De novo phased assembly of the Vitis riparia grape genome. Sci Data. 2019;6:127.

17. Xiao H, Liu Z, Wang N, Long Q, Cao S, Huang G, et al. Adaptive and maladaptive introgression in grapevine domestication. Proceedings of the National Academy of Sciences. 2023;120:e2222041120.

18. Zecca G, Labra M, Grassi F. Untangling the Evolution of American Wild Grapes: Admixed Species and How to Find Them. Front Plant Sci. 2020;10:1814.

19. Ma Z-Y, Wen J, Ickert-Bond SM, Nie Z-L, Chen L-Q, Liu X-Q. Phylogenomics, biogeography, and adaptive radiation of grapes. Mol Phylogenet Evol. 2018;129:258–67.

20. Ho SYW, Duchêne S. Molecular-clock methods for estimating evolutionary rates and timescales. Molecular Ecology. 2014;23:5947–65.

21. Garrison E, Guarracino A. Unbiased pangenome graphs. Bioinformatics. 2023;39:btac743.

22. Hickey G, Heller D, Monlong J, Sibbesen JA, Sirén J, Eizenga J, et al. Genotyping structural variants in pangenome graphs using the vg toolkit. Genome Biol. 2020;21:35.

23. Marçais G, Delcher AL, Phillippy AM, Coston R, Salzberg SL, Zimin A. MUMmer4: A fast and versatile genome alignment system. Darling AE, editor. PLoS Comput Biol. 2018;14:e1005944.

24. Cingolani P, Platts A, Wang LL, Coon M, Nguyen T, Wang L, et al. A program for annotating and predicting the effects of single nucleotide polymorphisms, SnpEff: SNPs in the genome of Drosophila melanogaster strain w ^1118^; iso-2; iso-3. Fly. 2012;6:80–92.

25. Morales-Cruz A, Aguirre-Liguori J, Massonnet M, Minio A, Zaccheo M, Cochetel N, et al. Multigenic resistance to Xylella fastidiosa in wild grapes (Vitis sps.) and its implications within a changing climate. Commun Biol. 2023;6:1–15.

26. Heinitz CC, Riaz S, Tenscher AC, Romero N, Walker MA. Survey of chloride exclusion in grape germplasm from the southwestern United States and Mexico. Crop Sci. 2020;60:1946–56.

27. Myles S, Boyko AR, Owens CL, Brown PJ, Grassi F, Aradhya MK, et al. Genetic structure and domestication history of the grape. Proceedings of the National Academy of Sciences. 2011;108:3530–5.

28. Tello J, Ibáñez J. Review: Status and prospects of association mapping in grapevine. Plant Science. 2023;327:111539.

29. Garg S, Balboa R, Kuja J. Chromosome-scale haplotype-resolved pangenomics. Trends in Genetics. 2022;38:1103–7.

30. Riaz S, Pap D, Uretsky J, Laucou V, Boursiquot J-M, Kocsis L, et al. Genetic diversity and parentage analysis of grape rootstocks. Theor Appl Genet. 2019;132:1847–60.

31. Li N, He Q, Wang J, Wang B, Zhao J, Huang S, et al. Super-pangenome analyses highlight genomic diversity and structural variation across wild and cultivated tomato species. Nat Genet. 2023;55:852–60.

32. Shang L, Li X, He H, Yuan Q, Song Y, Wei Z, et al. A super pan-genomic landscape of rice. Cell Res. 2022;32:878–96.

33. Tao Y, Luo H, Xu J, Cruickshank A, Zhao X, Teng F, et al. Extensive variation within the pan-genome of cultivated and wild sorghum. Nat Plants. 2021;7:766–73.

34. Tang D, Jia Y, Zhang J, Li H, Cheng L, Wang P, et al. Genome evolution and diversity of wild and cultivated potatoes. Nature. 2022;606:535–41.

35. Bennetzen JL, Wang H. The Contributions of Transposable Elements to the Structure, Function, and Evolution of Plant Genomes. Annual Review of Plant Biology. 2014;65:505–30.

36. Fedoroff N. Transposons and genome evolution in plants. Proc Natl Acad Sci U S A. 2000;97:7002–7.

37. Zou C, Massonnet M, Minio A, Patel S, Llaca V, Karn A, et al. Multiple independent recombinations led to hermaphroditism in grapevine. Proceedings of the National Academy of Sciences. 2021;118:e2023548118.

38. Charlesworth D, Charlesworth B, Marais G. Steps in the evolution of heteromorphic sex chromosomes. Heredity. 2005;95:118–28.

39. Golicz AA, Bayer PE, Bhalla PL, Batley J, Edwards D. Pangenomics Comes of Age: From Bacteria to Plant and Animal Applications. Trends in Genetics. 2020;36:132–45.

40. Vondras AM, Minio A, Blanco-Ulate B, Figueroa-Balderas R, Penn MA, Zhou Y, et al. The genomic diversification of grapevine clones. BMC Genomics. 2019;20:972.

41. Liu Y. A code within the genetic code: codon usage regulates co-translational protein folding. Cell Communication and Signaling. 2020;18:145.

42. Carrasco D, Zhou-Tsang A, Rodriguez-Izquierdo A, Ocete R, Revilla MA, Arroyo-García R. Coastal Wild Grapevine Accession (Vitis vinifera L. ssp. sylvestris) Shows Distinct Late and Early Transcriptome Changes under Salt Stress in Comparison to Commercial Rootstock Richter 110. Plants. 2022;11:2688.

43. Ning L, Kan G, Shao H, Yu D. Physiological and transcriptional responses to salt stress in salt-tolerant and salt-sensitive soybean (Glycine max [L.] Merr.) seedlings. Land Degradation & Development. 2018;29:2707–19.

44. Guan R, Qu Y, Guo Y, Yu L, Liu Y, Jiang J, et al. Salinity tolerance in soybean is modulated by natural variation in GmSALT3. The Plant Journal. 2014;80:937–50.

45. Lin J, Massonnet M, Cantu D. The genetic basis of grape and wine aroma. Hortic Res. 2019;6:1–24.

46. Chin C-S, Peluso P, Sedlazeck FJ, Nattestad M, Concepcion GT, Clum A, et al. Phased diploid genome assembly with single-molecule real-time sequencing. Nat Methods. 2016;13:1050–4.

47. Cochetel N, Minio A, Massonnet M, Vondras AM, Figueroa-Balderas R, Cantu D. Diploid chromosome-scale assembly of the *Muscadinia rotundifolia* genome supports chromosome fusion and disease resistance gene expansion during *Vitis* and *Muscadinia* divergence. Morrell P, editor. G3 Genes|Genomes|Genetics. 2021;11:jkab033.

48. Blanco-Ulate B, Vincenti E, Powell ALT, Cantu D. Tomato transcriptome and mutant analyses suggest a role for plant stress hormones in the interaction between fruit and Botrytis cinerea. Front Plant Sci. 2013;4:142.

49. Lam ET, Hastie A, Lin C, Ehrlich D, Das SK, Austin MD, et al. Genome mapping on nanochannel arrays for structural variation analysis and sequence assembly. Nat Biotechnol. 2012;30:771–6.

50. Waterhouse RM, Seppey M, Simão FA, Manni M, Ioannidis P, Klioutchnikov G, et al. BUSCO Applications from Quality Assessments to Gene Prediction and Phylogenomics. Molecular Biology and Evolution. 2018;35:543–8.

51. Wang M, Kong L. pblat: a multithread blat algorithm speeding up aligning sequences to genomes. BMC Bioinformatics. 2019;20:28.

52. Smit A, Hubley R, Green P. RepeatMasker Open-4.0. 2013.

53. Bolger AM, Lohse M, Usadel B. Trimmomatic: a flexible trimmer for Illumina sequence data. Bioinformatics. 2014;30:2114–20.

54. Andrews S. FastQC: A Quality Control tool for High Throughput Sequence Data. 2014.

55. Pertea M, Pertea GM, Antonescu CM, Chang T-C, Mendell JT, Salzberg SL. StringTie enables improved reconstruction of a transcriptome from RNA-seq reads. Nat Biotechnol. 2015;33:290– 5.

56. Grabherr MG, Haas BJ, Yassour M, Levin JZ, Thompson DA, Amit I, et al. Full-length transcriptome assembly from RNA-Seq data without a reference genome. Nat Biotechnol. 2011;29:644–52.

57. Au KF, Underwood JG, Lee L, Wong WH. Improving PacBio Long Read Accuracy by Short Read Alignment. Xing Y, editor. PLoS ONE. 2012;7:e46679.

58. Haas BJ. Improving the Arabidopsis genome annotation using maximal transcript alignment assemblies. Nucleic Acids Res. 2003;31:5654–66.

59. Slater G, Birney E. Automated generation of heuristics for biological sequence comparison. BMC Bioinformatics. 2005;6:31.

60. Boratyn GM, Thierry-Mieg J, Thierry-Mieg D, Busby B, Madden TL. Magic-BLAST, an accurate RNA-seq aligner for long and short reads. BMC Bioinformatics. 2019;20:405.

61. Stanke M, Keller O, Gunduz I, Hayes A, Waack S, Morgenstern B. AUGUSTUS: ab initio prediction of alternative transcripts. Nucleic Acids Res. 2006;34:W435–9.

62. Lomsadze A. Gene identification in novel eukaryotic genomes by self-training algorithm. Nucleic Acids Res. 2005;33:6494–506.

63. Korf I. Gene finding in novel genomes. BMC Bioinformatics. 2004;5:59.

64. Haas BJ, Salzberg SL, Zhu W, Pertea M, Allen JE, Orvis J, et al. Automated eukaryotic gene structure annotation using EVidenceModeler and the Program to Assemble Spliced Alignments. Genome Biol. 2008;9:R7.

65. Gotz S, Garcia-Gomez JM, Terol J, Williams TD, Nagaraj SH, Nueda MJ, et al. High-throughput functional annotation and data mining with the Blast2GO suite. Nucleic Acids Res. 2008;36:3420–35.

66. Camacho C, Coulouris G, Avagyan V, Ma N, Papadopoulos J, Bealer K, et al. BLAST+: architecture and applications. BMC Bioinformatics. 2009;10:421.

67. Buchfink B, Xie C, Huson DH. Fast and sensitive protein alignment using DIAMOND. Nat Methods. 2014;12:59.

68. Jones P, Binns D, Chang H-Y, Fraser M, Li W, McAnulla C, et al. InterProScan 5: genome-scale protein function classification. Bioinformatics. 2014;30:1236–40.

69. Wu TD, Watanabe CK. GMAP: a genomic mapping and alignment program for mRNA and EST sequences. Bioinformatics. 2005;21:1859–75.

70. El-Gebali S, Mistry J, Bateman A, Eddy SR, Luciani A, Potter SC, et al. The Pfam protein families database in 2019. Nucleic Acids Res. 2019;47:D427–32.

71. Steuernagel B, Witek K, Krattinger SG, Ramirez-Gonzalez RH, Schoonbeek H, Yu G, et al. The NLR-Annotator Tool Enables Annotation of the Intracellular Immune Receptor Repertoire. Plant Physiol. 2020;183:468–82.

72. Alexa A, Rahnenfuhrer J. topGO: Enrichment Analysis for Gene Ontology. 2016.

73. Doležel J, Sgorbati S, Lucretti S. Comparison of three DNA fluorochromes for flow cytometric estimation of nuclear DNA content in plants. Physiol Plant. 1992;85:625–31.

74. Doležel J, Bartoš J. Plant DNA Flow Cytometry and Estimation of Nuclear Genome Size. Ann Bot. 2005;95:99–110.

75. Bertier L, Leus L, D’hondt L, de Cock AWAM, Höfte M. Host Adaptation and Speciation through Hybridization and Polyploidy in Phytophthora. PLOS ONE. 2013;8:e85385.

76. Smith TW, Kron P, Martin SL. flowPloidy: An R package for genome size and ploidy assessment of flow cytometry data. Appl Plant Sci. 2018;6:e01164.

77. Lodhi MA, Reisch BI. Nuclear DNA content of Vitis species, cultivars, and other genera of the Vitaceae. Theoret Appl Genetics. 1995;90:11–6.

78. Cochetel N, Cantu D. A super-pangenome of the North American wild grape species. GitHub. https://github.com/noecochetel/North_American_Vitis_Pangenome (2023).

79. Cochetel N, Cantu D. A super-pangenome of the North American wild grape species. Zenodo. 10.5281/zenodo.10206475 (2023).

80. Wang Y, Tang H, DeBarry JD, Tan X, Li J, Wang X, et al. MCScanX: a toolkit for detection and evolutionary analysis of gene synteny and collinearity. Nucleic Acids Res. 2012;40:e49.

81. Patro R, Duggal G, Love MI, Irizarry RA, Kingsford C. Salmon provides fast and bias-aware quantification of transcript expression. Nat Methods. 2017;14:417–9.

82. Srivastava A, Malik L, Sarkar H, Zakeri M, Almodaresi F, Soneson C, et al. Alignment and mapping methodology influence transcript abundance estimation. Genome Biology. 2020;21:239.

83. Soneson C, Love MI, Robinson MD. Differential analyses for RNA-seq: transcript-level estimates improve gene-level inferences. F1000Research. 2016;4.

84. Cochetel N, Minio A, Guarracino A, Garcia JF, Figueroa-Balderas R, Massonnet M, et al. A super-pangenome of the North American wild grape species. PRJNA984685. The base-level super-pangenome graph of the North American wild grape species (Vitis spp.) unveiled genus wide associations. https://www.ncbi.nlm.nih.gov/bioproject/?term=PRJNA984685 (2023).

85. Morales-Cruz A, Aguirre-Liguori JA, Zhou Y, Minio A, Riaz S, Walker AM, et al. Introgression among North American wild grapes (Vitis) fuels biotic and abiotic adaptation. PRJNA731597. Introgression among North American wild grapes (Vitis) fuels biotic and abiotic adaptation. https://www.ncbi.nlm.nih.gov/bioproject/?term=PRJNA731597 (2023).

86. Morales-Cruz A, Aguirre-Liguori J, Massonnet M, Minio A, Zaccheo M, Cochetel N, et al. Multigenic resistance to Xylella fastidiosa in wild grapes (Vitis sps.) and its implications within a changing climate. PRJNA842753. Multigenic resistance to Xylella fastidiosa in wild grapes (Vitis sps.) and its implications within a changing climate. https://www.ncbi.nlm.nih.gov/bioproject/?term=PRJNA842753 (2023).

87. Li H, Durbin R. Fast and accurate short read alignment with Burrows-Wheeler transform. Bioinformatics. 2009;25:1754–60.

88. Danecek P, Bonfield JK, Liddle J, Marshall J, Ohan V, Pollard MO, et al. Twelve years of SAMtools and BCFtools. GigaScience. 2021;10:giab008.

89. Quinlan AR, Hall IM. BEDTools: a flexible suite of utilities for comparing genomic features. Bioinformatics. 2010;26:841–2.

90. Chang CC, Chow CC, Tellier LC, Vattikuti S, Purcell SM, Lee JJ. Second-generation PLINK: rising to the challenge of larger and richer datasets. GigaSci. 2015;4:7.

91. Emms DM, Kelly S. OrthoFinder: phylogenetic orthology inference for comparative genomics. Genome Biology. 2019;20:238.

92. Edgar RC. MUSCLE: multiple sequence alignment with high accuracy and high throughput. Nucleic Acids Res. 2004;32:1792–7.

93. Castresana J. Selection of Conserved Blocks from Multiple Alignments for Their Use in Phylogenetic Analysis. Molecular Biology and Evolution. 2000;17:540–52.

94. Darriba D, Posada D, Kozlov AM, Stamatakis A, Morel B, Flouri T. ModelTest-NG: A New and Scalable Tool for the Selection of DNA and Protein Evolutionary Models. Molecular Biology and Evolution. 2020;37:291–4.

95. Kozlov AM, Darriba D, Flouri T, Morel B, Stamatakis A. RAxML-NG: a fast, scalable and user-friendly tool for maximum likelihood phylogenetic inference. Bioinformatics. 2019;35:4453–5.

96. Suchard MA, Lemey P, Baele G, Ayres DL, Drummond AJ, Rambaut A. Bayesian phylogenetic and phylodynamic data integration using BEAST 1.10. Virus Evolution. 2018;4:vey016.

97. Wan Y, Schwaninger HR, Baldo AM, Labate JA, Zhong G-Y, Simon CJ. A phylogenetic analysis of the grape genus (Vitis L.) reveals broad reticulation and concurrent diversification during neogene and quaternary climate change. BMC Evol Biol. 2013;13:141.

98. Suyama M, Torrents D, Bork P. PAL2NAL: robust conversion of protein sequence alignments into the corresponding codon alignments. Nucleic Acids Research. 2006;34:W609–12.

99. Yang Z. PAML 4: Phylogenetic Analysis by Maximum Likelihood. Mol Biol Evol. 2007;24:1586–91.

100. Álvarez-Carretero S, Kapli P, Yang Z. Beginner’s Guide on the Use of PAML to Detect Positive Selection. Crandall K, editor. Molecular Biology and Evolution. 2023;40:msad041.

101. Enright AJ, Van Dongen S, Ouzounis CA. An efficient algorithm for large-scale detection of protein families. Nucleic Acids Research. 2002;30:1575–84.

102. Mendes FK, Vanderpool D, Fulton B, Hahn MW. CAFE 5 models variation in evolutionary rates among gene families. Bioinformatics. 2020;36:5516–8.

103. Hill WG, Weir BS. Variances and covariances of squared linkage disequilibria in finite populations. Theoretical Population Biology. 1988;33:54–78.

104. Nicolas SD, Péros J-P, Lacombe T, Launay A, Le Paslier M-C, Bérard A, et al. Genetic diversity, linkage disequilibrium and power of a large grapevine (Vitis vinifera L) diversity panel newly designed for association studies. BMC Plant Biol. 2016;16:74.

105. Zhou X, Stephens M. Genome-wide efficient mixed-model analysis for association studies. Nat Genet. 2012;44:821–4.

106. McKenna A, Hanna M, Banks E, Sivachenko A, Cibulskis K, Kernytsky A, et al. The Genome Analysis Toolkit: A MapReduce framework for analyzing next-generation DNA sequencing data. Genome Res. 2010;20:1297–303.

107. RStudio Team. RStudio: Integrated Development Environment for R [Internet]. Boston, MA: RStudio, Inc.; 2022. Available from: http://www.rstudio.com/

108. R Core Team. R: A language and environment for statistical computing. R Foundation for Statistical Computing, Vienna, Austria; 2022.

109. Wickham H, Averick M, Bryan J, Chang W, McGowan L, François R, et al. Welcome to the Tidyverse. Journal of Open Source Software. 2019;4:1686.

110. Lawrence M, Huber W, Pagès H, Aboyoun P, Carlson M, Gentleman R, et al. Software for Computing and Annotating Genomic Ranges. Prlic A, editor. PLoS Comput Biol. 2013;9:e1003118.

111. Yu G. Using ggtree to Visualize Data on Tree-Like Structures. Current Protocols in Bioinformatics. 2020;69:e96.

112. Wang L-G, Lam TT-Y, Xu S, Dai Z, Zhou L, Feng T, et al. Treeio: An R Package for Phylogenetic Tree Input and Output with Richly Annotated and Associated Data. Molecular Biology and Evolution. 2020;37:599–603.

113. Conway JR, Lex A, Gehlenborg N. UpSetR: an R package for the visualization of intersecting sets and their properties. Bioinformatics. 2017;33:2938–40.

114. Wilkins D. gggenes: Draw Gene Arrow Maps in “ggplot2” [Internet]. 2023. Available from: https://CRAN.R-project.org/package=gggenes

115. Gu Z, Gu L, Eils R, Schlesner M, Brors B. circlize implements and enhances circular visualization in R. Bioinformatics. 2014;30:2811–2.

116. Cochetel N, Minio A, Guarracino A, Garcia JF, Figueroa-Balderas R, Massonnet M, et al. A super-pangenome of the North American wild grape species. Zenodo. 10.5281/zenodo.8065808 (2023).

